# Concomitant post-translational repression of *Arabidopsis* PIP1 aquaporins upon the loss of major PIP2 isoforms

**DOI:** 10.64898/2026.08.14.744787

**Authors:** Komal Jhala, Jessica M. Lehnert, Birgit Geist, Juliane Merl-Pham, Jin Zhao, Chen Liu, Anton R. Schäffner

## Abstract

Aquaporins at the plant plasmalemma are divided into two highly conserved subclasses, PLASMA MEMBRANE INTINSIC PROTEINs 1 (PIP1) and PIP2. A*rabidopsis thaliana* encodes five PIP1 and eight PIP2 isoforms. Individual loss-of-function mutants had been employed for functional analyses. Here, we observe that the *pip2;1 pip2;2 pip2;4 pip2;6 pip2;7* quintuple mutant defective of major PIP2 isoforms concomitantly leads to a strongly reduced PIP1 protein level. Lower order mutants *pip2;1 pip2;2* and *pip2;1 pip2;2 pip2;7* still harbor only 60% and 20% residual PIP1, respectively. This repression is established post-translationally, since neither *PIP1*’s steady-state transcripts nor polysome-associated *PIP1* mRNAs are suppressed by *pip2;1 pip2;2 pip2;7*. Thus, the two major pathways operating in eukaryotes for removal of aberrant proteins, ubiquitin proteasome system (UPS)-dependent ER-associated degradation (ERAD) and autophagy/vacuole-linked degradation, were assessed. Introgression of *atg7* blocking autophagy-mediated degradation does not affect the PIP1 protein level of *pip2;1 pip2;2 pip2;7*. In contrast, introgression of ERAD loss-of-function mutations *hrd1A hrd1B* and *dln1* into *pip2;1 pip2;2 pip2;7* partially stabilizes its PIP1 protein level. PIP1 accumulates intracellularly upon pharmacological inhibition of proteasomal degradation by MG132. Nevertheless, the lack of a full PIP1 recovery by these means suggests the flexible operation of parallel ERAD components or unknown pathways. In conclusion, the essential dependence of PIP1 expression on PIP2 isoforms intrinsically interconnects the two PIP subclades at the protein level and will thereby affect their mutual functions.

**Significance statement:** Plasma membrane intrinsic proteins constituting the most homogenous plant aquaporin family are nonetheless split into two highly conserved subfamilies, PIP1 and PIP2. The loss of major *Arabidopsis* PIP2 isoforms does not lead to compensation by PIP1 members, but rather to PIP1s’ concomitant, post-translational repression. This dependence of PIP1 isoforms inevitably ties the two PIP subfamilies and their function.

## Introduction

Water acquisition and homeostasis are vital for the survival of any organism. Aquaporins are passive water channels of the MIP (major intrinsic protein) family. Certain isoforms not only allow the passage of water but of other small, uncharged molecules, *e.g*., CO_2_, NH_3,_ H_2_O_2_, and arsenic, boric, or silicic acid. PLASMA MEMBRANE INTRINSIC PROTEINs (PIPs) comprise the most abundant and highly conserved plant aquaporin family. They play crucial roles in plant cell osmoregulation, root and leaf hydraulic conductivity, transpiration, and cell elongation (Maurel et al, 2015; Afzal et al, 2016). The earliest evolutionary occurrence of a PIP is documented for the colony-forming alga *Coccomyxa*. A split into two paralogous groups, PIP1 and PIP2, with highly conserved amino acid signatures is first documented in mosses and absolutely conserved in all land plants (Kammerloher et al, 1994; Schäffner, 1998; Alexandersson et al, 2005; Danielson and Johanson, 2008; Anderberg et al, 2011; Soto et al, 2012; Abascal et al, 2014; Maurel et al, 2015; Bienert et al, 2018). This suggests a crucial requirement for both PIP groups and/or their cooperative interaction. Plant genomes harbor more than 10 *PIP* isoforms, with usually more *PIP2* than *PIP1* members; *Arabidopsis thaliana* encodes five PIP1 and eight PIP2 proteins among a total of 35 MIP members (Yaneff et al, 2014; Maurel et al, 2015). Nearly 70% of the total PIP protein amount of 21-day-old *Arabidopsis* rosettes is dominated by PIP1;2 and PIP2;1 followed by PIP1;1 and PIP2;7 contributing about 10%. The remaining 20% are constituted by PIP1;1, PIP1;3, PIP1;5, PIP2;3, and PIP2;6. In 49-day-old roots, 70% of the total PIP protein is dominated by PIP1;1, PIP1;2, PIP2;1, and PIP2;2 followed by PIP2;4, PIP2;7, and PIP1;5 (Monneuse et al, 2011).

PIP monomers constitute functional channels formed by two half-helices each containing a central, conserved NPA signature sequence; however, their function requires a tetrameric assembly (Törnroth-Horsefield et al, 2006). This assembly provides an important feature of the interaction of PIP1 and PIP2, since PIP isoforms do also associate into PIP1-PIP2 heterotetrameric entities which enhance the water permeability of the participating PIP proteins and alter gating properties of the complex. Another important feature of PIP1-PIP2 heteromerization is the guidance of PIP1 isoforms to the plasma membrane (PM) which are otherwise retained at the ER membrane and show suppressed PM localization and, consequently, low water permeability (Fetter et al, 2004; Zelazny et al, 2007; Otto et al, 2010; Bienert et al, 2012; Chevalier et al, 2014; Yaneff et al, 2014; Jozefkowicz et al, 2016; Bienert et al, 2018). The latter observation originally led to underestimating the water channel activity of PIP1 proteins, which were frequently classified as non-active. However, in other instances, PIP1-PIP2 pairs did not functionally interact, and individual PIP1 were also functional water channels even in heterologous systems (Kammerloher et al, 1994; Fetter et al, 2004; Zhou et al, 2007; Horie et al, 2011; Jozefkowicz et al, 2013).

Studies employing loss-of-function mutants revealed several physiological and developmental impacts of PIP aquaporins. E.g., PIP2;1 is enhancing root-shoot water relocation, it facilitates the emergence of lateral roots, and it may be involved in stomatal function due to its water- and H_2_O_2_- transporting property (Da Ines et al, 2010; Péret et al, 2012; Grondin et al, 2015; Wang et al, 2016; Rodrigues et al, 2017; Ceciliato et al, 2019). PIP1;4 was shown to be involved in H_2_O_2_ signalling in response to bacterial pathogens (Tian et al, 2016).

Plant aquaporins are regulated at the transcriptional and post-translational level, *e.g*., in response to environmental cues (Maurel et al, 2021). Drought or salinity downregulates the transcript levels of many *PIP* genes; this was interpreted to reduce the loss of water from the plant to the environment (Jang et al, 2004; Alexandersson et al, 2005). Post-translational regulation includes PIP1-PIP2 tetramerization (Zelazny et al, 2007; Bienert et al, 2012; Yaneff et al, 2014; Jozefkowicz et al, 2016) and protein phosphorylation to control the gating of aquaporins by shifting open *vs*. closed states (Prado et al, 2013; Grondin et al, 2015; Rodrigues et al, 2017) or their subcellular trafficking (Dhonukshe et al, 2007; Prak et al, 2008; Boursiac et al, 2008; Wudick et al, 2015; Pou et al, 2016; Ueda et al, 2016). The controlled degradation of aquaporins at the PM or enroute to the PM involving elements of the ERAD pathway and autophagy, is a significant aspect of plant responses to environmental stimuli (Lee et al, 2009; Hachez et al, 2014a; Chen et al, 2021; Chen et al, 2022). AtPIP2;1 was shown to be polyubiquitinated by the pepper ubiquitin ligase Rma1H1 (RING membrane-anchor1), followed by degradation via the proteasome (Lee et al, 2009). Chen et al (2021) showed that the ERAD components, E2 enzyme UBC32 (ubiquitin-conjugating enzyme) and E3 ligase enzyme RMA1, act in concert as E2-E3 pair to positively regulate drought tolerance in *Arabidopsis thaliana* by targeting PIP2;1 and PIP2;2 for degradation. In other studies, AtPIP2;1 was endocytosed either via clathrin-coated vesicles under resting conditions (Dhonukshe et al, 2007) or in a raft-associated manner in response to salt stress (Li et al, 2011). SNARE proteins and aquaporins can interact with each other impacting water permeation through aquaporins (Hachez et al, 2014b; Baena et al, 2024). These different pathways and physiological situations leading to the targeting, retention or degradation of PIP proteins suggest the existence of multiple layers of regulation fine tuning the PIP protein levels at the PM.

The ER-associated degradation (ERAD) is part of the ubiquitin proteasome system (UPS) which is a conserved quality control mechanism across eukaryotes (Mehrtash and Hochstrasser, 2019). ERAD employs distinct sets of membrane-localized multiprotein complexes to identify unfolded, misfolded or unassembled proteins distinguishing the location of the defect within the ER membrane (ERAD-M), in the ER lumen (ERAD-L), or in the cytosol (ERAD-C) (Vembar and Brodsky, 2008). The central component of these multiprotein complexes is an E3 ubiquitin ligase exposed to the cytosolic surface of the ER membrane, which not only ubiquitinates ERAD substrates but also connects to various ER luminal and cytosolic adapters (Kumari and Brodsky, 2021). Most of our knowledge of ERAD stems from extensive research in yeast. Yeast contains at least two distinct E3 ligases, Hrd1 (HMG-CoA reductase degradation) and Doa10 (Degradation of alpha2) that ubiquitinate ERAD-L/M and ERAD-C substrates, respectively. There are two genes coding for Hrd1 homologs of Arabidopsis, *AtHRD1A* and *AtHRD1B* (Su et al, 2011). Emerging evidence from Arabidopsis suggests that plants preserve the conserved Hrd1-centered ERAD machinery, while simultaneously evolving additional components that modulate its activity in accordance with plant-specific physiological contexts (Wu et al, 2026).

In addition, yeast Dfm1 (Derlin1-like Family Member 1) takes part in ERAD-M as part of the Hrd1 complex as well as in ERAD-C associated with the Doa10 complex. Yeast Dfm1 has been implicated in the degradation of membrane proteins (Neal et al, 2018). It acts along with the Hrd1 complex to recruit Cdc48 to the membrane. Dfm1 also promotes Hrd1-independent retrotranslocation of membrane proteins by supporting the Doa10 pathway (Christianson and Carvalho, 2022). It can also function autonomously and participates in the retrotranslocation of integral membrane substrates of Hrd1 and Doa10 pathway with the aid of Cdc48 (Neal et al, 2018, 2020). Additionally, Dfm1 may exhibit chaperone-like activity (Kandel *et al,* 2023). There are three genes encoding DERLINs in *Arabidopsis*, *AtDERLIN1* (*AtDLN1*), *AtDLN2A*, and *AtDLN2B*. *AtDLN1* is the orthologue of yeast Dfm1 as revealed through bioinformatic analysis (Paul et al, 2013). Dfm1 has homologous counterparts across plants, yeast, and animals, but its role in plants remains elusive.

Autophagy is another degradation pathway conserved across eukaryotes, initially discovered in yeast as well (Xie and Klionsky, 2007). Most autophagy-related (*ATG*) genes are well conserved throughout evolution, and their homologs exist in mammals and plants (Yamamoto et al, 2023). The yeast mutants defective in Atg5 and Atg7 showed accumulation of defective membrane proteins (Lipatova *et al,* 2020). The important role of Atg5 and Atg7 has also been extended to mammalian cells (Molinari *et al,* 2021). Autophagic degradation of PIP aquaporins has been implicated in the response of *Arabidopsis* to drought. AtPIP2;7 was shown to interact with a TSPO membrane protein to form a complex that was degraded through autophagy (Hachez et al, 2014a). In *Medicago truncatula* ATG8 binding to dehydrin MtCAS31 facilitated the autophagic degradation of MtPIP2;7 that eventually improved drought resistance of the plants (Li et al, 2020). In mammalian systems, the degradation of AQP2 under hypokalemic conditions has been shown to depend on ATG7, based on studies using Atg7-deficient mice (Kim et al, 2019).

Here, we report that loss of major *Arabidopsis* PIP2 isoforms PIP2;1, PIP2;2, and PIP2;7 leads to a concomitant repression of all PIP1 proteins instead of initiating a compensatory response by inducing PIP1 proteins. Tagged PIP1 proteins confirmed the reduction of the respective PIP1 isoform in *pip2* mutant backgrounds *in planta* and demonstrated the impact of PIP2 proteins on PIP1s’ targeting and stability when transiently expressed in protoplasts. Transcriptional and translational analyses indicated that the repression of PIP1 happened at the post-translational level. Reverse genetic experiments and pharmacological inhibition suggest that the degradation of PIP1 proteins in the absence of PIP2;1, PIP2;2, and PIP2;7 was at least partially ERAD-associated. Thus, the absence of these major PIP2 isoforms affected both the localization and abundance of PIP1s, revealing a scenario controlling the interplay of PIP aquaporins in plant cells.

## Results

### The loss of major *PIP2* isoforms leads to a concomitant downregulation of PIP1 proteins

Plant plasma membrane aquaporins are abundant proteins and split into two subclades, PIP1 and PIP2 (Robinson et al, 1996; Schäffner, 1998; Monneuse et al, 2011; Maurel et al, 2015). To comprehensively study the function of the major PIP2 isoforms of *Arabidopsis thaliana*, a multiple loss-of-function mutant *pip2;1 pip2;2 pip2;4 pip2;6 pip2;7* (quintuple *pip2* mutant) was established which would eliminate 99% of PIP2 proteins in leaves and roots (Monneuse et al, 2011). The expression profiles of *PIP2;1, PIP2;2, PIP2;4, PIP2;6*, and *PIP2;7* examined by transgenic *PIP2_pro_::GUS* reporter lines and by proteomic analyses showed that *PIP2;1*, *PIP2;2*, and *PIP2;7* are highly expressed in roots as well as in leaves. *PIP2;4* is root-specific and strongly expressed in lateral and young roots but not in the hypocotyl, whereas *PIP2;6* is predominantly detected in young leaves and the main root (Jang et al, 2004; Da Ines, 2008; Monneuse et al, 2011).

*pip2;1 pip2;2 pip2;4 pip2;6 pip2;7* was viable and grew only somewhat smaller under regular cultivation (Figure S1). We reasoned that the loss of these major PIP2 isoforms might have an impact on the PIP1-type aquaporins. Therefore, microsomal fractions of wild type and the quintuple *pip2* mutant rosette leaves were probed by ELISA using an anti-PIP1 antiserum detecting all five *Arabidopsis* PIP1 isoforms (Henzler et al, 1999; Figure S2). Notably, the loss of the PIP2 isoforms was concomitant with a strong reduction of PIP1 proteins by 90% instead of any potential compensatory effect (Figure 1a). Lower-order *pip2* mutants *pip2;1 pip2;2*, *pip2;1 pip2;2 pip2;7*, and *pip2;1 pip2;2 pip2;4 pip2;6* were analyzed to identify the contribution of individual *PIP2* genes. The *pip2;1 pip2;2* double mutant showed a reduction of PIP1 by about 30%, the *pip2;1 pip2;2 pip2;7* triple mutant by nearly 70%, and *pip2;1 pip2;2 pip2;4 pip2;6* by 50% (Figure 1a). Thus, the introgression of *pip2;7* into *pip2;1 pip2;2* or into *pip2;1 pip2;2 pip2;4 pip2;6* mutants further reduced the PIP1 protein level by about 40%. The PIP1 protein level of *pip2;1 pip2;2* and *pip2;1 pip2;2 pip2;7* was independently addressed by Western blotting (Figure 1b). In agreement with the previous result, there was a significant decrease in PIP1 protein by 80% in *pip2;1 pip2;2 pip2;7* and by 50% in the *pip2;1 pip2;2* compared to wild-type levels (Figure 1c). Thus, all studied PIP2 isoforms contribute to the gradually stronger repression of PIP1 protein level, culminating in the extreme downregulation observed by the *pip2* quintuple mutant. Moreover, it appears that PIP2;1, PIP2;2, and PIP2;7 are the isoforms primarily affecting PIP1 levels in accordance with their dominant contribution to PIP2 proteins in rosette leaves (94% according to Monneuse et al, 2011).

**Figure 1.**
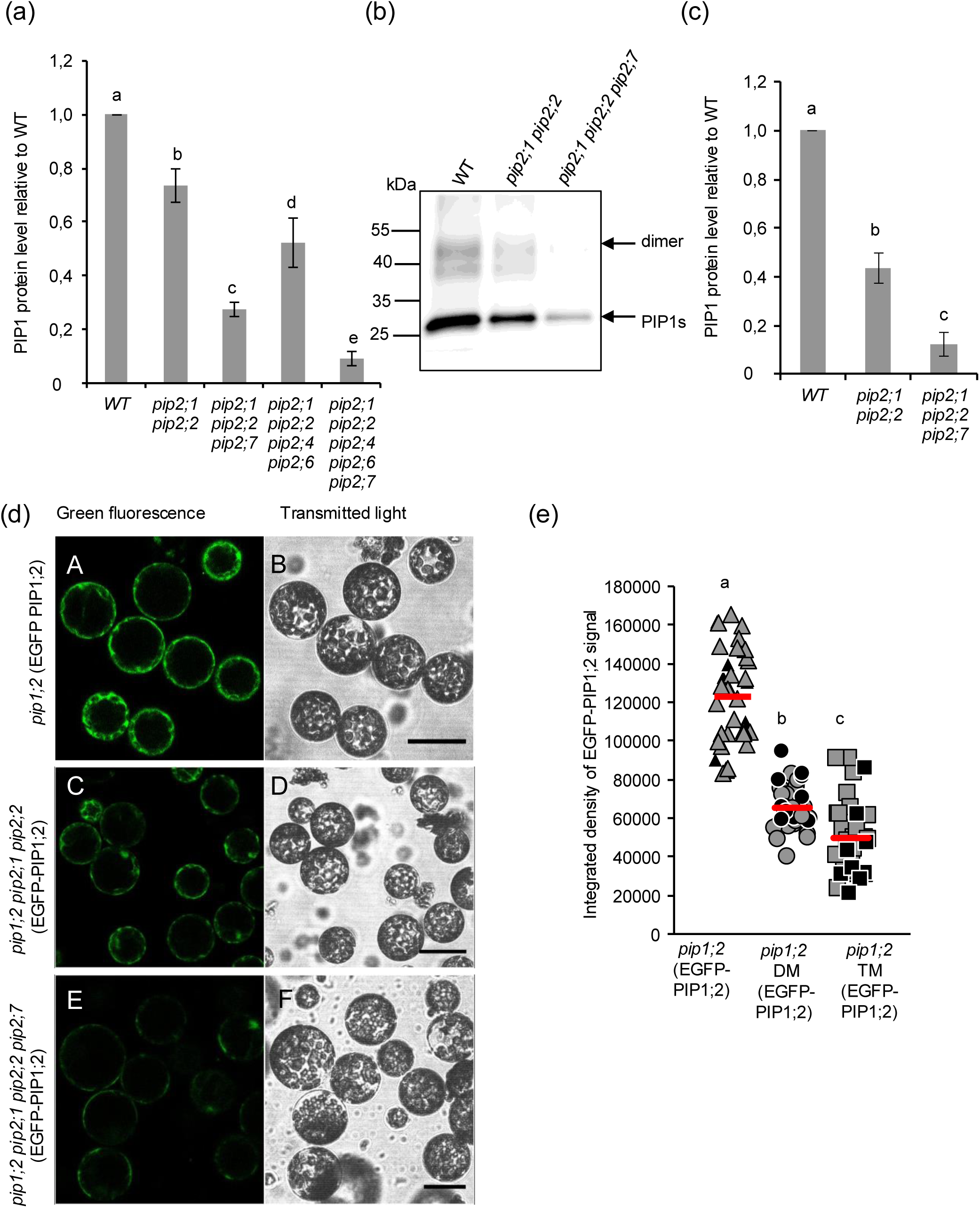
PIP1 proteins are downregulated in higher order *pip2* mutants. (a) ELISA assay of higher order *pip2* mutants. Microsomal fractions were isolated from 14- day-old rosette leaves. Polyclonal anti-PIP1 antiserum detecting all five PIP1 isoforms was used to probe PIP1 (Suppl. Fig. 2). Bars indicate means ± SD of three independent experiments. One-way ANOVA with Tukey’s post hoc method was used to examine significant differences (P< 0.050) in pairwise comparison and classified by letters. WT, wild type. (b) Western blot analysis of *pip2* mutants with polyclonal anti-PIP1 antiserum (monomers at approximately 28 kDa, dimers at 52 kDa). Parallelly run Coomassie-stained gels of the same, denatured protein sample demonstrated equal protein loading (Figure S3a). (c) PIP1 protein level was quantified from Western blots using Image J and normalized to the corresponding lanes of a parallel, Coomassie-stained SDS polyacrylamide gel. The protein level was finally normalized to WT. Bars are means ± SD of three independent experiments. One-way ANOVA with Tukey’s post hoc method was used to examine significant differences (P< 0.001) in pairwise comparison and classified by letters. WT, wild type. (d) Mesophyll protoplasts were isolated from one-month-old plants of *pip1;2* (EGFP-PIP1;2), *pip1;2* DM (EGFP-PIP 1;2) and *pip1;2* TM (EGFP-PIP 1;2) plants grown on soil with a 10-h-light/14-h-dark cycle. EGFP-PIP1;2 fluorescence was measured by confocal microscopy. Representative images are shown. Scale bar, 50 µm. DM*: pip2;1 pip2;2* and TM*: pip2;1 pip2;2 pip2;7*. (e) Integrated density was calculated using Image J software (Fiji) for 36 protoplasts of each line. Data are represented as scatter plot with the red horizontal line defining the mean value. Kruskal-Wallis one-way ANOVA with the Tukey’s post hoc method was performed to examine significant differences (P < 0.050) in pairwise comparison classified by letters. The integrated density of the protoplasts shown in (d) are denoted in the graph by black color for the corresponding genotype. The integrated density for some of the protoplasts in the representative images is overlapping and is therefore not visible in the graph.

### All five PIP1 protein isoforms are repressed in the *pip2;1 pip2;2* mutant

The anti-PIP1 antiserum globally assesses the PIP1 members. To explore which specific PIP1 isoforms were repressed, *pip2;1 pip2;2* double mutant microsomal proteins were compared to wild type employing a proteomic approach. *pip2;1 pip2;2* was chosen for this study as it still contains about half of the PIP1 protein pool, thus enabling the quantification of residual PIP1 isoforms. Microsomal membrane fractions from 28-day-old leaves and 21-day-old roots of wild type and of *pip2;1 pip2,2* were subjected to LC-MS/MS-based label-free peptide quantification. All five PIP1 isoforms were identified by mass spectrometry in root samples, whereas all PIP1s except PIP1;1 were detected in leaf microsomal fractions. A general and significant reduction of all PIP1 isoforms was revealed in *pip2;1 pip2;2* compared to wild-type samples. Only about 20 to 60% normalized peptide peak intensities of individual PIP1 proteins were detected, *i.e.*, all PIP1 isoforms were strongly affected (Table 1).

**Table 1.** All five PIP1 isoforms are repressed in the *pip2;1 pip2;2* double mutant compared to the wild type based on LC-MS/MS-based label-free quantification.

| Tissue | Name | AGI code | Unique peptides | Fold change | Raw p-value | Adjusted p-value |
| --- | --- | --- | --- | --- | --- | --- |
| **Rosette** | PIP1;2 | AT2G45960 | 1 | 0.40 | 8.46E-07 | 0.000239 |
|  | PIP1;3 | AT1G01620 | 2 | 0.53 | 9.41E-05 | 0.005277 |
|  | PIP1;4 | AT4G00430 | 2 | 0.58 | 0.000466 | 0.013773 |
|  | PIP1;5 | AT4G23400 | 2 | 0.36 | 4.54E-12 | 1.15E-08 |
| **Root** | PIP1;1 | AT3G61430 | 2 | 0.18 | 1.20E-07 | 8.43E-05 |
|  | PIP1;2 | AT2G45960 | 3 | 0.34 | 9.04E-09 | 2.39E-05 |
|  | PIP1;3 | AT1G01620 | 3 | 0.50 | 0.000112 | 0.008486 |
|  | PIP1;4 | AT4G00430 | 3 | 0.57 | 0.00107 | 0.023566 |
|  | PIP1;5 | AT4G23400 | 3 | 0.21 | 1.70E-08 | 2.39E-05 |

### Fluorescence-tagged PIP1 is downregulated in *pip2* mutant lines

In addition to PIP1 protein quantification, the level of a single, major PIP1 isoform was independently examined *in vivo* by employing transgenic mutant lines expressing EGFP-PIP1;2 under the control of its native promoter (*PIP1;2_pro_*::*EGFP-PIP1;2*) in the *pip1;2* single mutant background, *pip1;2 pip2;1 pip2;2*, and *pip1;2 pip2;1 pip2;2 pip2;7*. Thereby, the introgressed labeled *PIP1;2* isoform established a pseudo PIP1;2 wild-type situation in the lines harboring the *pip1;2* loss-of-function allele. Confocal microscopy was employed to quantify the EGFP fluorescence of mesophyll protoplasts isolated from *pip1;2* (EGFP-PIP1;2), *pip1;2 pip2;1 pip2;2* (EGFP-PIP1;2), and *pip1;2 pip2;1 pip2;2 pip2;7* (EGFP-PIP1;2) transgenic plants. EGFP-PIP1;2 fluorescence was significantly reduced to 48% and 61% in *pip1;2 pip2;1 pip2;2* (EGFP-PIP1;2) and *pip1;2 pip2;1 pip2;2 pip2;7* (EGFP-PIP1;2), respectively, compared to *pip1;2* (EGFP-PIP1;2) (Figure 1d, e).

### PM targeting and amount of PIP1;2 protein is enhanced by PIP2;1

To analyze whether the loss of PIP2s also affects the targeting of PIP1 proteins in addition to their quantity PIP1;2-EGFP was transiently expressed in *pip1;2 pip2;1 pip2;2* mesophyll protoplasts alone or together with PIP2;1 employing *35S_pro_::PIP1;2-EGFP* and *d35S_pro_::HA-PIP2;1* plasmid constructs. The PIP1;2-EGFP signal was observed by confocal microscopy. The exclusive expression of PIP1;2- EGFP exhibited weak and patchy signals at the PM, whereas a strong PIP1;2-EGFP signal at the PM was observed when PIP1;2-EGFP and HA-PIP2;1 had been coexpressed (Figure 2a, Figure S4).

**Figure 2.**
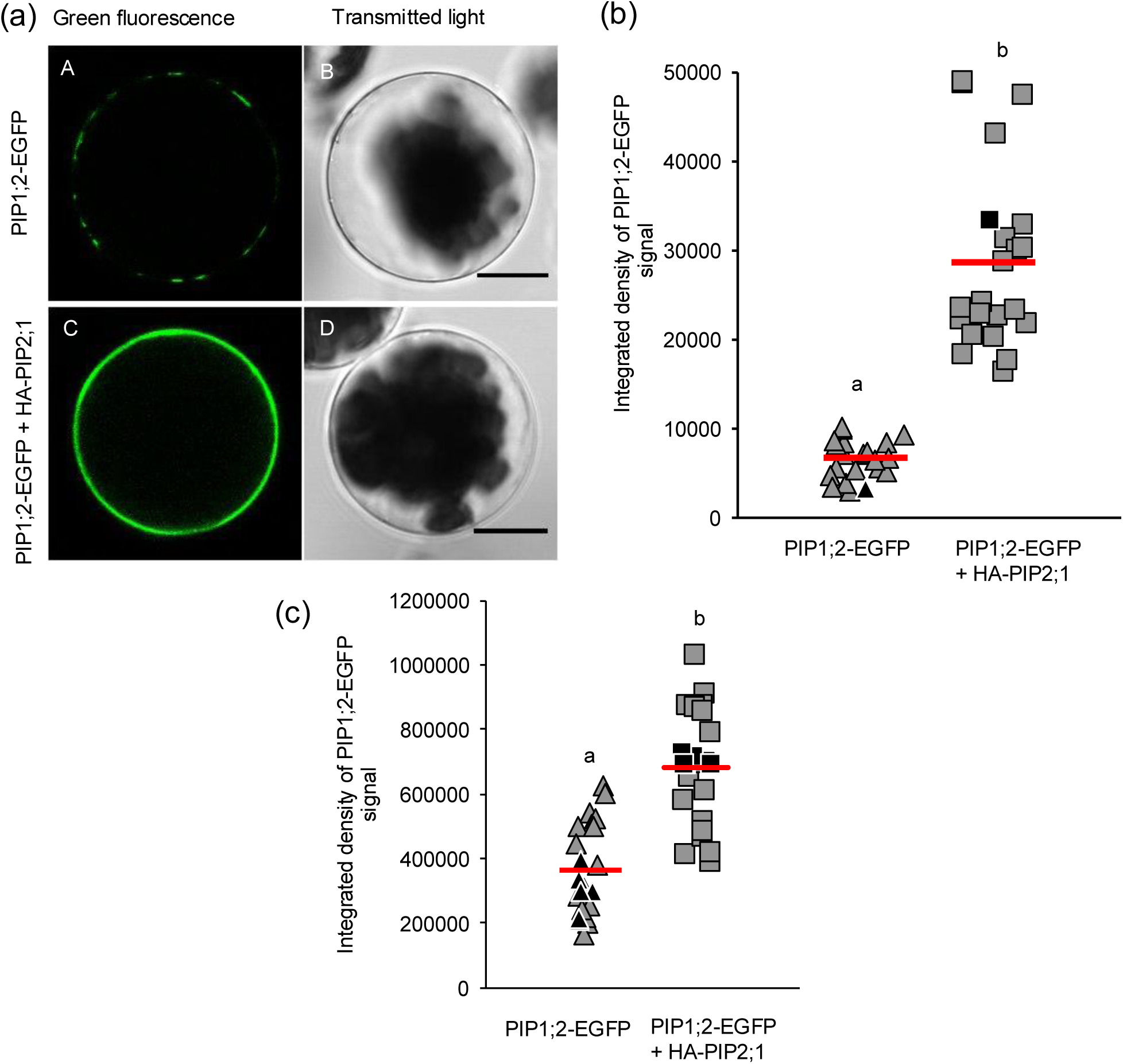
Targeting and abundance of PIP1;2 is influenced by PIP2;1. (a) *pip1;2 pip2;1 pip2;2* mesophyll protoplast were transfected with *35S_pro_*::*PIP1;2-EGFP* alone or co-transfected together with 2x*35S_pro_*::*HA-PIP2;1*. EGFP fluorescence was monitored 16 h after transfection. The upper panels (A, B) depict a cross section of a protoplast expressing PIP1;2- EGFP alone, whereas the lower panels (C, D) show the co-expression of PIP1;2-EGFP with PIP2;1. Scale bar, 50 µm. (b) Quantification of PIP1;2-EGFP fluorescence signal at the plasma membrane from central cross sections of the protoplasts to assess the targeting of PIP1;2. Data are plotted as integrated density of the PIP1;2-EGFP signal from protoplasts expressing PIP1;2-EGFP alone and together with HA- PIP2;1 (n = 22 protoplasts). Data are represented as scatter plot with the red horizontal line defining the mean value. Mann-Whitney Rank sum test was performed to examine significant differences (P<0.001) classified by letters. The integrated density of the protoplasts shown in Fig. 2a are denoted in the graph by black color for the corresponding condition. (c) Quantification of total PIP1;2-EGFP fluorescence signal from transfected protoplast. Example images of protoplasts used for quantification are shown in Fig. S4. Data are plotted as integrated density of the PIP1;2-EGFP signal from protoplasts expressing PIP1;2-EGFP alone and together with HA-PIP2;1 (n = 22 protoplasts). Data are represented as scatter plot with the red horizontal line defining the mean value. t-test was performed to examine significant differences (P<0.001) classified by letters. The integrated density of the protoplasts shown in Fig. S4 are denoted in the graph by black color for the corresponding condition.

We also quantified the total fluorescence of PIP1;2-EGFP affected by PIP2;1 apart from PIP1;2 PM targeting. Therefore, EGFP fluorescence signal intensity was collected for protoplasts expressing PIP1;2-EGFP alone or together with PIP2;1. Interestingly, the PIP1;2 signal doubled when co- expressed with PIP2;1 (Figure 2b, Figure S4). This suggests that PIP2;1 supports PIP1 expression and assists PIP1s to reach the PM.

### PIP1;1 and PIP1;2 physically interact with PIP2 isoforms

According to the histochemical localization of *PIP1* and *PIP2* expression using promoter-reporter lines as well as cell-type specific transcriptomics, certain *PIP1* and *PIP2* genes exhibit overlapping expression patterns, which is an essential prerequisite for a mutual impact or physical interaction between members of these two subfamilies (Birnbaum et al, 2003; Javot et al, 2003; Da Ines, 2008; Alexandersson et al, 2010; Da Ines et al, 2010; Postaire et al, 2010; Péret et al, 2012; Prado et al, 2013; https://bar.utoronto.ca). The dependence of the PIP1 protein level on PIP2;1, PIP2;2, and PIP2;7 could involve a physical interaction of these proteins. To assess whether PIP1 and PIP2 isoforms of *Arabidopsis* interact, microsomal membrane fractions from transgenic lines *pip1;1 (PIP1;1_pro_::HA-PIP1;1)* and *pip1;1 pip2;1 pip2;2 (PIP1;1_pro_::HA-PIP1;1)* were subjected to a co- immunoprecipitation analysis. HA-tagged PIP1;1 proteins were immunoprecipitated from these microsomal fractions with an anti-HA antibody and probed by Western blotting using an anti-PIP2 antiserum recognizing the highly related PIP2;1, PIP2;2, and PIP2;3 (Da Ines, 2008). PIP2 proteins were co-precipitated from the microsomal fraction of the HA-PIP1;1-complemented pseudo wild- type line (Figure 3a). In contrast, the PIP2;1/PIP2;2/PIP2;3 signal was not detected from *pip1;1 pip2;1 pip2;2* (*HA-PIP1;1*), or from the HA-PIP1;1-complemented pseudo wild-type line when it had been processed without the anti-HA antibody (Figure 3a). Equivalent results were obtained with microsomal fractions from *pip1;2* (*PIP1;2_pro_::HA-PIP1;2*) and *pip1;2 pip2;1 pip2;2* (*PIP1;2_pro_::HA- PIP1;2*) (Figure S5).

**Figure 3.**
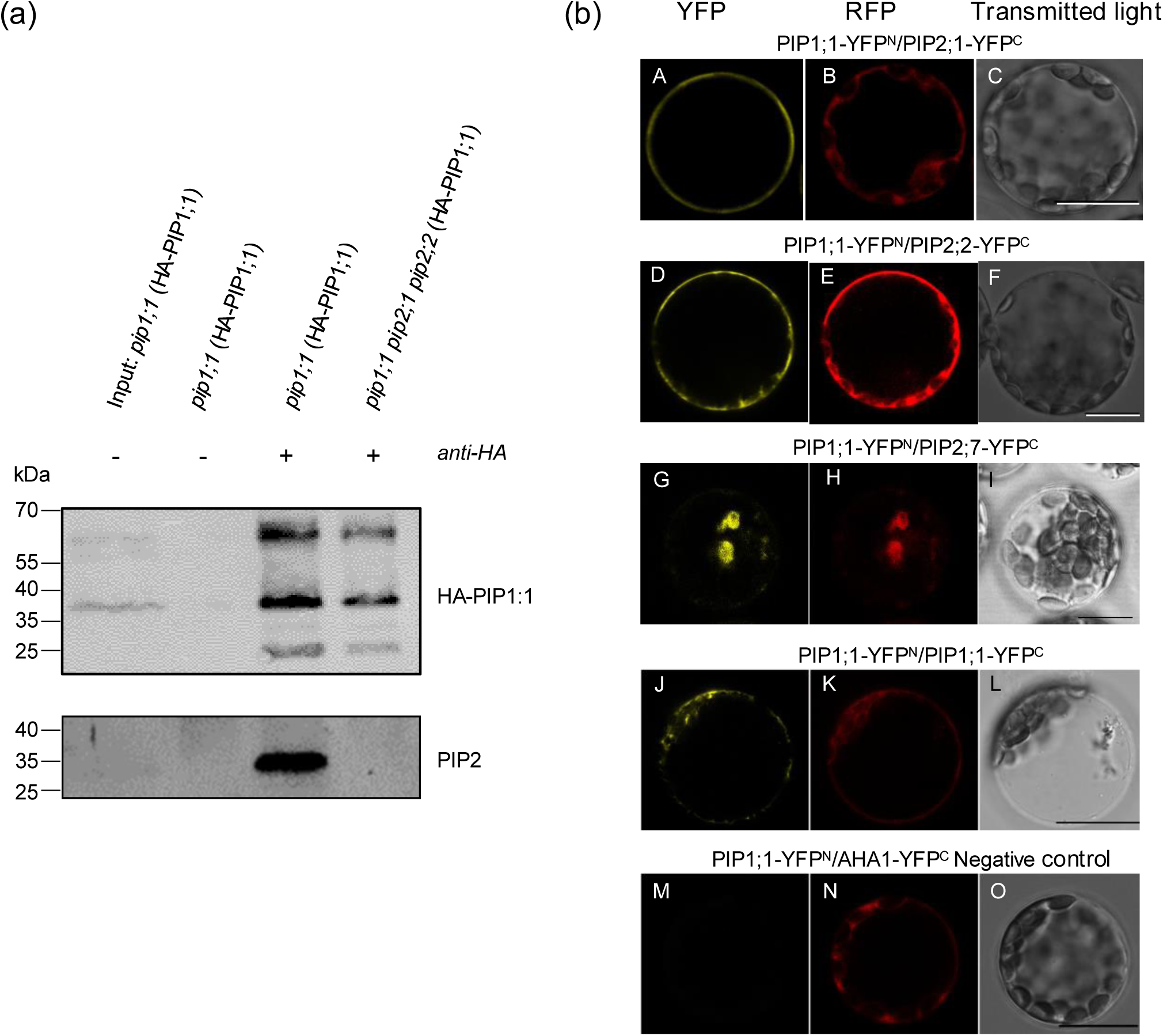
PIP1 and PIP2 proteins interact with each other impacting the targeting of PIP1. (a) Microsomal membrane fractions extracted from 28-day-old rosette leaves of transgenic *pip1;1* (HA-PIP1;1) and *pip1;1 pip2;1 pip2;2* (HA-PIP1;1) were subjected to immunoprecipitation using an anti-HA antibody and analyzed by Western blotting using the anti-HA antibody (upper panel, detecting HA-PIP1;1 monomer and dimer at about 36 kDa and 65 kDa) or an anti-PIP2;1/PIP2;2/PIP2;3 antiserum (lower panel, detecting approx. 28 kDa PIP2 monomer) (Materials and Methods). +, anti-HA antibody was added; – antibody was absent for immunoprecipitation. (b) Mesophyll protoplasts were isolated from *pip2;1 pip2;2 pip2;7* mutants were transformed with the BiFC constructs PIP1;1-YFP^N^/PIP2;1-YFP^C^, PIP1;1-YFP^N^/PIP2;2-YFP^C^, PIP1;1- YFP^N^/PIP2;7-YFP^C^, PIP1;1-YFP^N^/PIP1;1-YFP^C^ and PIP1;1-YFP^N^/AHA1-YFP^C^ (negative control). The constructs were visualized by confocal microscopy to observe YFP signal indicating protein-protein interaction (A, D, G, J, M), RFP signal as transformation control (B, E, H, K, N) and transmitted light (C, F, I, L, O). Representative images are shown. Scale bar, 50 μm.

Due to the high sequence similarity, the immunodetection could not decipher which PIP2 isoform interacts with PIP1;1 or PIP1;2. To pinpoint specific PIP2 isoforms, the elution fraction from the line expressing HA-PIP1;1 was further examined by LC-MS/MS. Peptides indicating the co- immunoprecipitation of PIP2;1, PIP2;2, PIP2;6, and PIP2;7 were identified (Table 2). Thus, other PIP2 isoforms except PIP2;1 and PIP2;2 may also interact with PIP1;1.

**Table 2.**
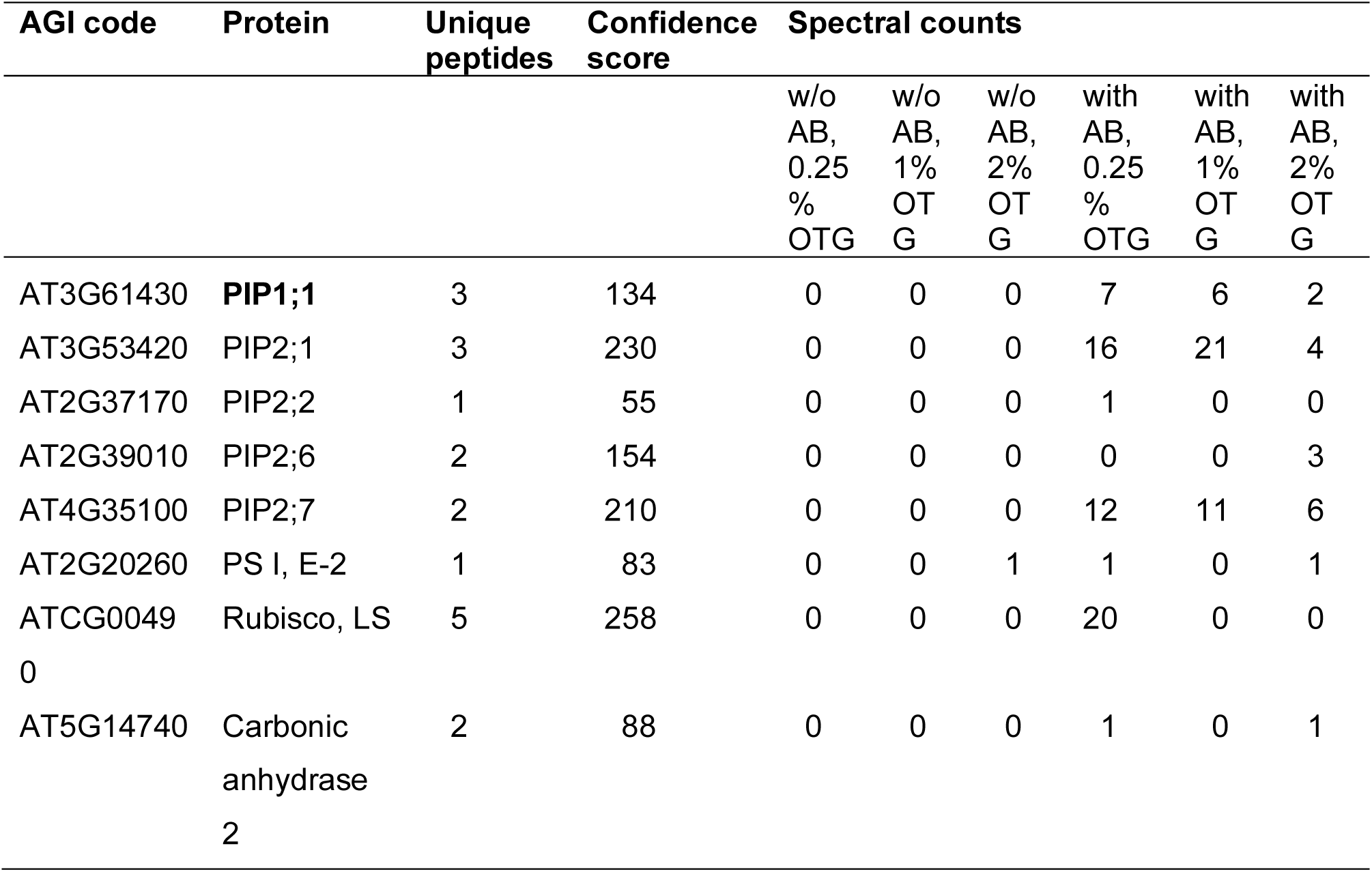
LC-MS/MS analysis of PIP1;1-HA pulldowns. After interaction with anti-HA antibodies and capture by protein A-agarose, the beads were washed with n-octyl-β-D-thioglucopyranoside (OTG)-containing buffers. Bound proteins were eluted with Laemmli buffer and examined by LC-MS/MS. Spectral counts indicated PIP2;1 and PIP2:7 as main interactors. However, PIP2;2 and PIP2;6 peptides were also identified in the elution fraction from the line expressing HA-PIP1;1. The spectral counts declined with increasing stringency of the washing (increasing OTG concentrations). The bait protein PIP1;1 is labeled in bold.

To further validate the physical interaction between the co-eluted proteins, bimolecular fluorescence complementation (BiFC) was employed. The vector pBiFCt-2in1-NN was used to generate constructs encoding chimeric fusions in which the potential PIP interactants were fused N-terminally to either the N-terminal (YFP^N^) or the C-terminal half (YFP^C^) of YFP (Grefen and Blatt, 2012). The chimeric constructs were transiently expressed in *pip2;1 pip2;2 pip2;7* mesophyll protoplasts; this mutant background was selected to avoid interference of major, endogenous PIP2s (Monneuse et al, 2011). The reconstitution of the YFP signal was assessed by confocal microscopy imaging. The co-transformed RFP signal confirms transformation also in the case of a missing interaction, i.e., in the absence of a YFP signal.

YFP signal at the PM was observed for PIP1;1-YFP^N^/PIP2;1-YFP^C^ and PIP1;1-YFP^N^/PIP2;2-YFP^C^ (Figure 3b) indicating the PIP1-PIP2 interaction and routing to the PM. In contrast, YFP signals were detected in intracellular structures and hardly at the PM for the pairs of PIP1;1-YFP^N^/PIP1;1-YFP^C^ and PIP1;1-YFP^N^/PIP2;7-YFP^C^ (Figure 3b). Thus, there is physical interaction between these pairs, but this interaction is not substantially contributing to PIP1;1’s routing to the PM. PIP1;1-YFP^N^/AHA1- YFP^C^ was chosen as negative control, since both the H^+^-ATPase AHA1 and PIP1;1 share PM localization, yet they are not physically interacting. Accordingly, no YFP signal was observed. In summary, these results demonstrated that PIP2;1, PIP2;2, and PIP2;7 physically interact with PIP1;1.

### *PIP1* genes are transcribed and translated irrespective of the presence of PIP 2;1, PIP 2;2, and PIP 2;7

To evaluate possible mechanisms behind the repression of PIP1 proteins in *pip2* mutants, all five *PIP1* genes were investigated by quantitative real-time PCR analysis to assess whether their transcriptional levels were altered in *pip2;1 pip2;2 pip2;7* mutants as compared to the wild type. RT- qPCR analyses using RNA from 21-day-old rosette leaves of WT and *pip2;1 pip2;2 pip2;7* showed that all five *PIP1* genes were transcribed in both genetic backgrounds, although *PIP1;1* and *PIP1;5* showed a downregulation by about 30-40% (Figure 4a). However, PIP1;1 and PIP1;5 only account for about 28% of the total PIP1 leaf proteome (Monneuse et al, 2011). Thus, the repression of PIP1 proteins is likely not linked to a reduced level of steady-state *PIP1* transcript levels.

**Figure 4.**
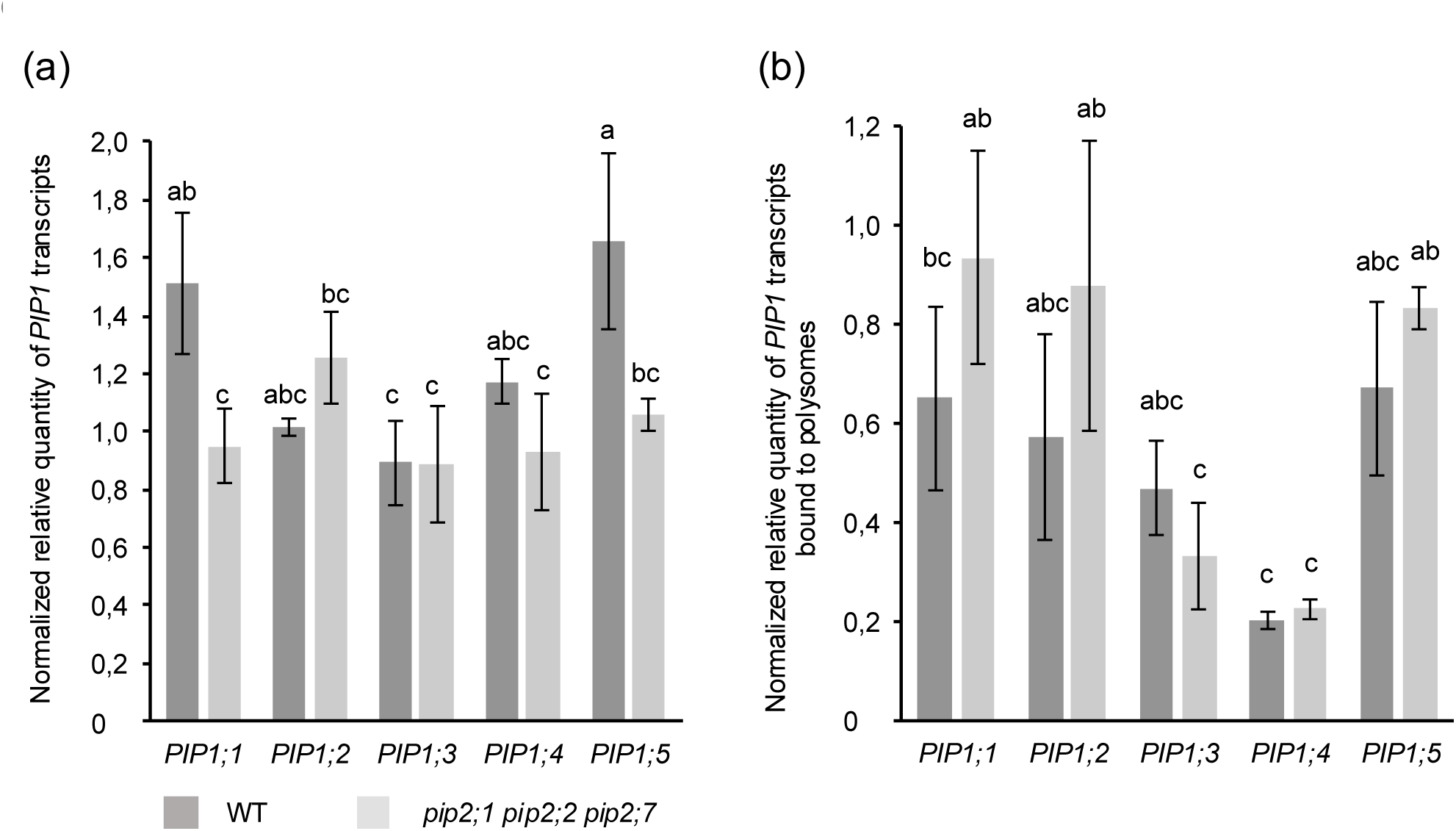
*PIP1* transcript abundance bound to polysomes does not change in the *pip2;1 pip2;2 pip2;7* mutant. (a) RNA was isolated from 21-day-old rosettes of wild type (dark bars) and *pip2;1 pip2;2 pip2;7* mutant (light grey bars). RT-qPCR data were normalized using stably expressed reference genes (Materials and Methods). Means ± SD of three independent experiments are displayed. One-way ANOVA with Tukey’s post hoc method was used to examine significant differences (P < 0.050) in pairwise comparison and classified by letters. WT. wild type. (b) RNA was isolated from the polysome fractions of the 21-day-old rosettes of the same genotypes as above. RT-qPCR data were normalized using stably expressed reference genes (Materials and Methods). Bars indicate means ± SD of three independent experiments. One-way ANOVA with Tukey’s post hoc method was used to examine significant differences (P< 0.050) in pairwise comparison and classified by letters. WT, wild type.

Therefore, we analyzed PIP1 expression at the translational level. Actively translated mRNAs are associated with multiple ribosomes in large polyribosome (polysome) complexes, whereas other mRNAs may remain as ribonucleoprotein complexes to be either stored or degraded (Proud, 2007). Accordingly, polysomes were isolated from *pip2;1 pip2;2 pip2;7* and wild type to assess the actual translation of *PIP1* mRNA. RT-qPCR analysis of the RNA extracted from polysomes did not reveal significant difference in transcript abundance between the wild type and the mutant (Figure 4b). Thus, PIP1 proteins are still translated in *pip2;1 pip2;2 pip2;7* like in wild type, despite the strong reduction in steady-state PIP1 protein level in the *pip2* triple mutant, arguing for a post-translational regulation.

### ERAD-mediated PIP1 degradation contributes to PIP1 repression in the absence of PIP2s

The active translation of PIP1, yet the eventual decrease in PIP1 at the steady state protein level suggests post-translational degradation in *pip2;1 pip2;2 pip2;7*. ERAD may be a potential mechanism involved in this regulation. Therefore, an interaction of PIP1 proteins with ERAD- associated UPS components at the ER, the E3 ligases HRD1B and RMA1, the adaptor protein HRD3A, the AAA-type ATPase CDC48A operating in the cytosol, and DLN1 was assessed by BiFC assays. The BiFC constructs PIP1;2-YFP^N^/DLN1-YFP^C^, PIP1;2-YFP^N^/HRD3A-YFP^C^, PIP1;2- YFP^N^/CDC48A-YFP^C^, PIP1;2-YFP^N^/RMA1-YFP^C^, PIP1;2-YFP^N^/HRD1B-YFP^C^, and HRD3A- YFP^N^/DLN1-YFP^C^ were established to test the interaction of the respective protein combinations. PIP1;2-YFP^N^/HRD1B-YFP^C^ and HRD3A-YFP^N^/DLN1-YFP^C^ were conceived as negative controls. HRD1B is an E3 ligase which does not directly interact with the faulty protein; therefore, PIP1;2 and HRD1B may not interact. HRD3A and DLN1 may constitute another negative control pair inferred from the yeast Hrd1 complex (ERAD-M) where Hrd3A and Dfm1 (yeast ortholog of AtDLN1) are part of the complex but do not physically interact (Mehrtash and Hochstrasser, 2019). The BiFC constructs were transiently expressed in *Arabidopsis pip1;2* mesophyll protoplasts to avoid interference with endogenous PIP1;2. Interestingly, reconstitution of the YFP signal indicating protein-protein interactions was observed for PIP1;2-DLN1, PIP1;2-HRD3A, and PIP1;2-CDC48A by confocal microscopy (Figure 5a). The PIP1;2-RMA1 pair and the conceived negative scenarios did not show interaction (Figure 5a).

**Figure 5.**
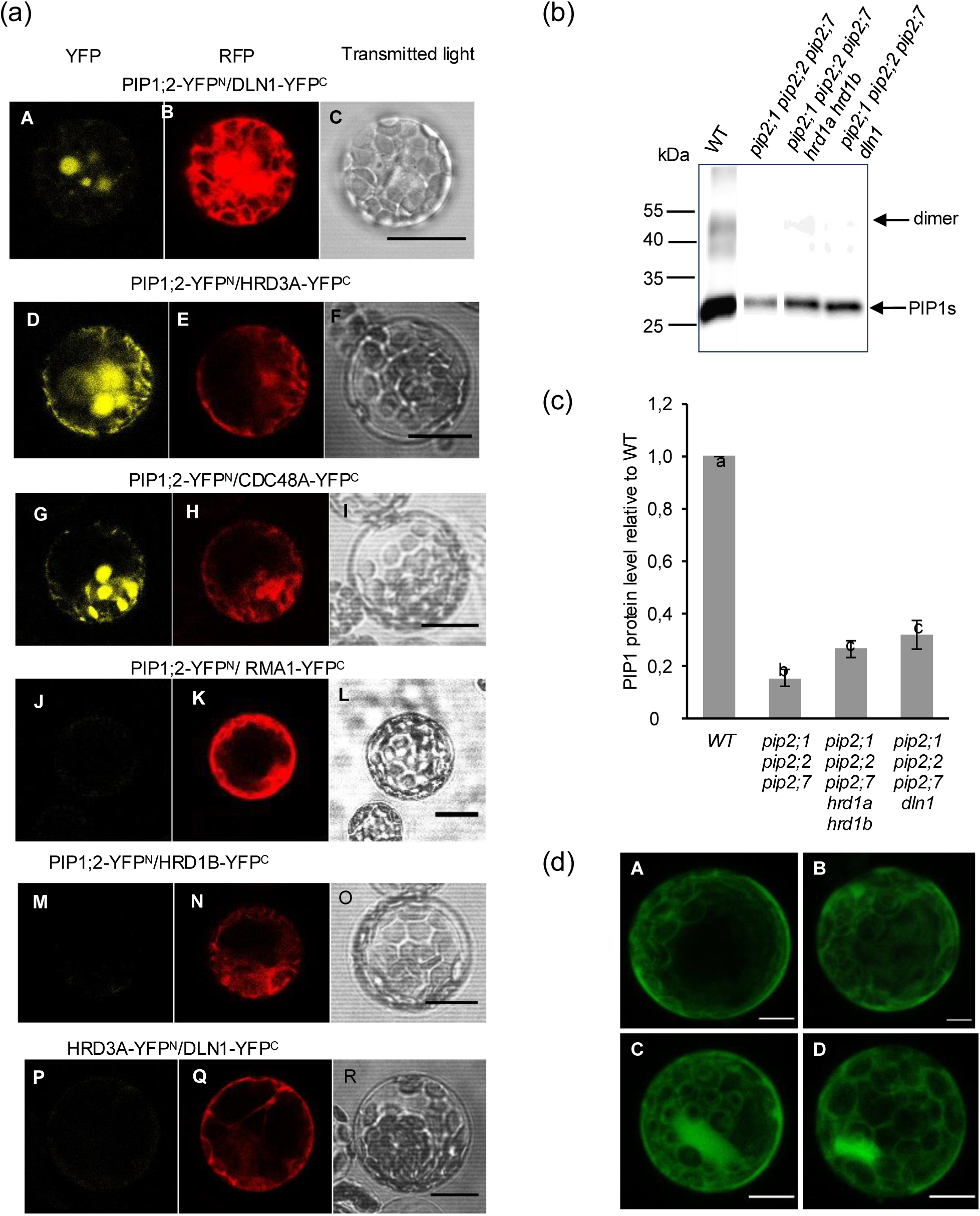
ERAD is partially involved in repressing PIP1 protein levels in absence of PIP2;1, PIP2;2, and PIP2;7. (a) *pip1;2* mesophyll protoplasts were transfected with the BiFC constructs PIP1;2-YFP^N^/DLN1- YFP^C^,PIP1;2 YFP^N^/HRD3A-YFP^C^, PIP1;2-YFP^N^/CDC48A-YFP^C^, PIP1;2-YFP^N^/RMA1-YFP^C^, PIP1;2-YFP^N^/HRD1B-YFP^C^ and HRD3A-YFP^N^/DLN1-YFP^C^. The PIP1;2-YFP^N^/HRD1B-YFP^C^ and HRD3A-YFP^N^/DLN1-YFP^C^ are predicted not to interact based on the known function of the homologous yeast ERAD components. The constructs were visualized by confocal microscopy to observe YFP signal indicating interaction (A, D, G, J, M, P), RFP signal as transfection control (B, E, H, K, N, Q), and transmitted light (C, F, I, L, O, R). Representative images are shown. Scale bar, 50 μm. (b) Microsomal fractions were isolated from 14-day-old rosette tissue of plants grown on plates. Polyclonal anti-PIP1 antiserum detecting all five PIP1 isoforms was used to probe PIP1 proteins (approximately 28 kDa monomers, 52 kDa dimers). (c) PIP1 protein level was quantified using image J by normalizing to the corresponding lane from a parallel, Coomassie-stained SDS polyacrylamide gel of the same denatured sample (Fig. S3b). The protein level was finally normalized to WT. Bars are means ± SD of three independent experiments. One way ANOVA with Tukey’s post hoc method was used to examine significant differences (P< 0.050) in pairwise comparison and classified by letters. WT, wild type (d) Mesophyll protoplasts were isolated from four-week-old *pip1;2 pip2;1 pip2;2 pip2;7* (EGFP- PIP1;2) and treated for 18 h with DMSO or 50 μM MG132. EGFP-PIP1;2 fluorescence was measured by confocal microscopy. Twelve to fourteen images of each protoplast were taken at Z intervals of 3 µm. Image J software (Fiji) was used to create sum of slice projections of the Z-stack images of the protoplasts. Representative images are shown (n = 15) (Figure S8). The protoplasts show a strong cytoplasmic accumulation of EGFP-PIP1;2 signal with MG132 treatment (C, D) compared to DMSO control (A, B). Scale bar, 50 μm.

The evidence of physical interaction of HRD3A, DLN1, and CDC48A with PIP1;2, supports a role of the HRD1 complex and therefore of ERAD in PIP1 regulation. We further hypothesized that the potential ER retention of PIP1, in an assembly defective state may trigger UPR (Unfolded Protein Response) and, thereby, it could be related to the observed PIP1 degradation. UPR signaling can be triggered by the accumulation of unfolded proteins at the ER (Howell, 2021). To assess UPR induction, the expression of *BIP3* and *BIP1/*2 genes which encode for chaperones assisting in protein folding and of *HRD1B* and *DLN1* which encode for proteins associated with the ERAD complex was assessed by RT-qPCR using RNA extracted from 14-day-old rosette leaves of WT and *pip2;1 pip2;2 pip2;7*. There was a twofold upregulation of *BIP1/2* and a non-significant, but consistent trend of slightly enhanced *BIP3*, *HRD1B*, and *DLN1* transcript levels (Figure S6). Thus, ERAD and UPR may indeed be affected in the *pip2;1 pip2;2 pip2;7* mutant.

Therefore, we further approached the role of HRD1A, HRD1B, and DLN1 in contributing to PIP1 regulation in the absence of PIP2 proteins by a reverse genetics approach. Two ERAD knockout mutants were generated in the *pip2;1 pip2;2 pip2;7* mutant background. The first mutant was generated by introgression of *hrd1a hrd1b*, a second one incorporated a *dln1* loss-of-function allele. The elimination of the ERAD-associated proteins might diminish the degradation of PIP1 leading to an accumulation of PIP1 proteins in the ERAD-defective *pip2;1 pip2;2 pip2;7* mutants. Microsomal fractions were isolated from 14-day-old rosette tissue of the WT and *pip2;1 pip2;2 pip2;7* mutants along with the set of ERAD-defective *pip2;1 pip2;2 pip2;7* mutants. PIP1 proteins were quantified by Western blotting using a polyclonal anti-PIP1 antiserum.

The ERAD-defective *pip2;1 pip2;2 pip2;7* mutants showed an about twofold increase in PIP1 protein level for both the *hrd1a hrd1b* and the *dln1* knockout situation. Thus, the stabilization of PIP1 is only partial and minor when compared to wild-type level (Figure 5b, c; Figure S7). In addition, we could observe intracellular accumulation of EGFP-PIP1;2 signals in mesophyll protoplasts isolated from the transgenic line *pip1;2 pip2;1 pip2;2 pip2;7* (*EGFP-PIP1;2*) when treated with the proteasome inhibitor MG132 (50 μM for 18 h) (Figure 5d; Figure S8a). Altogether these data show that ERAD is involved in downregulating PIP1 proteins in the absence of PIP2;1, PIP2;2, and PIP2;7.

### Autophagy does not play a sole role in PIP1 degradation

The disruption of ERAD has only a partial effect on the *pip2;1 pip2;2 pip2;7*-dependent PIP1 degradation. However, the UPS-dependent ERAD may act in parallel with the autophagy pathway to sustain protein homeostasis, and both pathways may complement each other (Minina et al, 2017). A crosstalk between the two pathways is well known in mammalian and yeast cells and for few instances from plant systems (Kocaturk and Gozuacik, 2018; Cao et al, 2021). Loss of Atg7 in yeast leads to the accumulation of overexpressed transmembrane proteins under normal growth conditions (Lipatova et al, 2020). Therefore, we hypothesized a parallel operation of autophagy along with ERAD in PIP1 turnover in the absence of PIP2;1, PIP2;2, and PIP2;7. Mutation of the *ATG7* gene results in autophagy deficiency, providing an ideal tool to study this degradation pathway (Doelling et al, 2002; Thompson et al, 2005). Therefore, the *pip2;1 pip2;2 pip2;7 atg7* mutant combination was generated and analyzed for accumulation of PIP1 proteins. However, *pip2;1 pip2;2 pip2;7 atg7* did not change the PIP1 protein level when compared with the *pip2;1 pip2;2 pip2;7* mutant (Figure 6a, b). Accordingly, ATG protein-mediated autophagy is not playing a crucial, at least sole role in PIP1 degradation in the absence of PIP2;1, PIP2;2 and PIP2;7.

**Figure 6.**
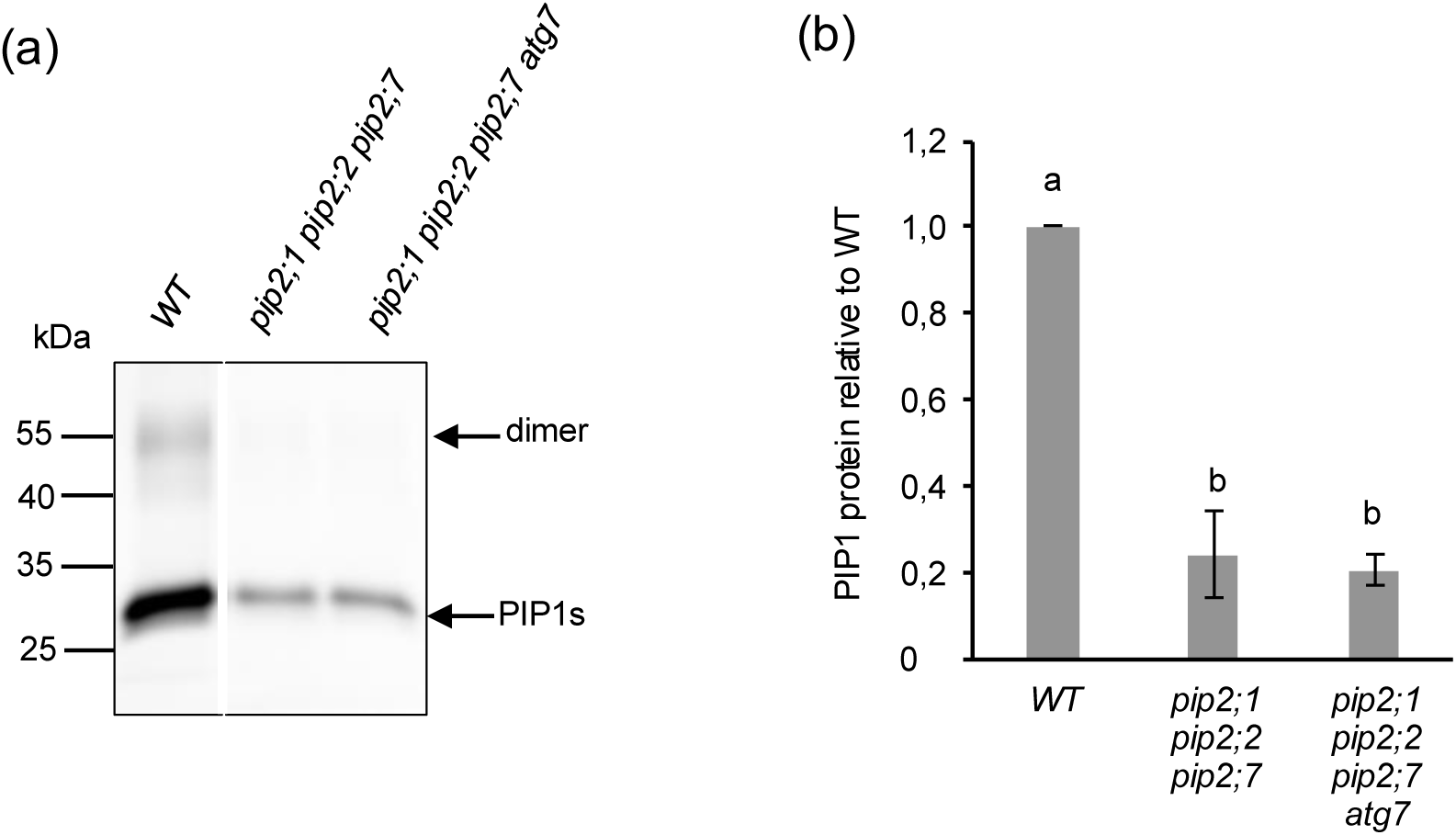
Defective autophagy is not recovering PIP1 protein levels in the absence of PIP2;1, PIP2;2, and PIP2;7. (a) Microsomal fractions were isolated from 14-day-old rosette leaves of plants grown on agar plates. Polyclonal anti-PIP1 antiserum detecting all five PIP1 isoforms was used to probe PIP1 proteins (approx. 28 kDa monomer, 52 kDa dimer). A parallelly run gel of the same, denatured protein samples was stained by Coomassie to demonstrate equal protein loading (Figure S3c). (b) PIP1 protein level was quantified from Western blots using Image J by normalizing to corresponding lanes of a parallel Coomassie-stained SDS polyacrylamide gel. The protein level was finally referred to WT. Bars are means ± SD of three independent experiments. One way ANOVA with Tukey’s post hoc method was used to examine significant differences (P< 0.050) in pairwise comparison and classified by letters. WT, wild type.

## Discussion

### PIP1-PIP2 interaction and implications

Plants are unique as they possess the highest number of PM-localized aquaporin isoforms among eukaryotes. PIPs appeared already in algae and are highly conserved in all higher plants. In addition, there is a strikingly strict conservation of their split into two paralogous subfamilies PIP1 and PIP2 which has been already observed in an ancestral embryophyte (Schäffner, 2008; Danielson and Johanson, 2008; Anderson et al, 2011; Abascal et al, 2014; Bienert et al, 2018). Accordingly, their interaction and relationship have been a crucial part of the study of PIP aquaporins. Numerous physical interactions of specific PIP1-PIP2 pairs have been documented which were basically substantiated by the structural assembly of PIP monomers into obligate tetramers allowing the formation of heterotetramers (Fetter et al, 2004; Temmei et al, 2005; Zelazny et al, 2007, 2009; Mahdieh et al, 2008; Matsumoto et al, 2009; Vandeleur et al, 2009; Bellati et al, 2010, 2016; Otto et al, 2010; Ayadi et al, 2011; Horie et al, 2011; Sorieul et al, 2011; Bienert et al, 2012; Jozefkowicz et al, 2013; Jones et al, 2014; Chevalier et al, 2014; Yaneff et al, 2014; Bienert et al, 2018; Shibasaka et al, 2021; Paluch-Lubawa and Polcyn, 2024). These interactions have been linked to a variety of PIP functions and regulations. Importantly, the interactions enabled the trafficking of the respective PIP1 isoforms to the PM whereas they were stuck in internal membranes without co-expressed, specific PIP2 proteins. Furthermore, these cooperative mechanisms of heterotetramer formation established latent or modified PIP functions. Thereby, the properties of PIPs were modulated leading to enhanced or interaction-dependent water permeability, to altered regulatory sensitivity, or to the regulation of PIP-related CO_2_ and ion conductance (Fetter et al, 2004; Temmei et al, 2005; Matsumoto et al, 2009; Bellati et al, 2010; Otto et al, 2010; Yaneff et al, 2014; Jozefkowicz et al, 2016; Byrt et al, 2017; Shibasaka et al, 2021; Tyerman et al, 2021). Furthermore, the multiplicity of PIPs is reflected in the interaction of several PIP2 isoforms with a particular PIP1 in planta (Table 2; Figure 3a, b; Figure S5). Since many other membrane proteins have been identified in protein-protein interactions with an *Arabidopsis* PIP1 and a PIP2 member (Belatti et al, 2016), the functionality of PIPs and their (hetero-)tetramers may even further multiply and diversify. On the other hand, the identified interactions were frequently isoform-specific, while other pairs failed to target the PIP1 partner to the PM or to affect PIP functions (Fetter et al, 2004; Secchi and Zwieniecki, 2010; Horie et al, 2011; Jozefkowicz et al, 2013). In case of overexpression experiments, like the interaction of PIP1;1 and PIP2;7 in this study (Figure 3b), the failure of PM targeting may be also attributed to a deficiency of endogenous proteins mediating the post-Golgi trafficking (Hachez et al, 2014b).

Here, we show an additional, principal level of the PIP1-PIP2 interaction, the expression and stability of PIP1 proteins fundamentally depending on the presence of PIP2 isoforms. Notably, the loss of PIP2 members does not lead to a compensatory effect but rather to the concomitant posttranslational downregulation and degradation of all PIP1 isoforms. Thereby, Arabidopsis PIP2;1, PIP2;2, and PIP2;7 play the predominant role in stabilizing PIP1 isoforms. However, the downregulation of PIP1s in absence of these PIP2s may not only be a matter of specificity, but rather related to their overlapping expression pattern as shown by histochemical analyses (Da Ines, 2008; Javot et al, 2003; Alexandersson et al, 2010; Da Ines et al, 2010; Postaire et al, 2010; Peret et al, 2012; Prado et al, 2013; Zhao, 2013) and, importantly, to the shear abundance of PIP2;1, PIP2;2, and PIP2;7 that make up 94% of PIP2 proteins in *Arabidopsis* leaves (Monneuse et al, 2011). Still, the additional loss of PIP2;4 and PIP2;6 had a further impact on the repression of PIP1 proteins (Figure 1).

### PIP1 downregulation occurs at the ER

*PIP1* genes are transcribed and translated in *pip2* loss-of-function mutants (Figure 4a, b), however, the post-translational processing is impaired, and the PIP1 protein level is reduced. The reduction of PIP1 proteins was analyzed using the microsomal fractions implying that they were indeed degraded instead of being mistargeted to any other membranes in *pip2* mutants. Since remaining PIP1 proteins accumulate at the ER (Figure 3b; Figure S4; Zelazny et al, 2007), we hypothesized that this accumulation activates ER quality control mechanisms, particularly degradation via ERAD or autophagy. PIP1;2 interacted with ERAD components, including the E3 ligases HRD1 and DLN1 and the AAA ATPase CDC48, implicating these proteins in PIP1 downregulation (Figure 5a). The introgression of *hrd1a hrd1b* or *dln1* into the *pip2;1 pip2;2 pip2;7* triple mutant was to severely compromise ERAD, however, led only to a partial PIP1 recovery (Figure 5b, c). In contrast, the blockage of autophagy in an *atg7*-deficient mutant background did not affect the PIP1 downregulation (Figure 6a, b). Thus, our biochemical and genetic studies revealed evidence for the partial involvement of ERAD, but not HRD1- or DLN1-dependent ERAD as a unique or principal degradation pathway. The modest increase in PIP1 protein level in the *pip2;1 pip2;2 pip2;7 hrd1a hrd1b* mutant hints to a probable activation of UBC32 protein which is otherwise controlled by the HRD1 protein in Arabidopsis (Chen et al, 2016). UBC32 is a Doa10 complex component in yeast. In Arabidopsis, UBC32, together with RMA1, ubiquitinates PIP2;1 and PIP2;2 under drought stress (Chen et al, 2021). Yeast Hrd3 remodelled itself to accommodate Dfm1 substrates, in particular, ERAD-M substrates in the dfm1 mutant strain (Neal et al, 2020). By analogy, HRD1 in Arabidopsis may possess the capacity to reorganize its substrate recognition or interaction landscape under conditions where canonical cofactors or client proteins are absent. We speculate that in the *pip2;1 pip2;2 pip2;7 dln1* mutant, HRD1 or its associated machinery may undergo functional “rebranding” to process non-canonical substrates such as unassembled PIP1.

An overlapping and parallel operation of ERAD components is documented in *Arabidopsis* wherein a single defective protein can undergo degradation via multiple ERAD component linked routes. The degradation of bri1-5 and bri1-9 mutant proteins was slowed down evident from their upregulation in the respective ERAD mutants, such as *hrd3a*, *hrd1a hrd1b* (Su *et al,* 2011), *ubc32* (Cui *et al,* 2012), *mns4 mns5* (Hüttner *et al,* 2014), and *ebs7* (Liu *et al,* 2015). Collectively, these observations suggest that although HRD1A/HRD1B and DLN1 likely play a role in mediating PIP1 degradation, their functional loss may be compensated by other ERAD components, including both characterized factors and potentially undiscovered elements of the ER quality control system. Furthermore, inhibition of the 26S proteasome using MG132 resulted in an enhanced intracellular accumulation of PIP1 proteins (Figure 5d), thereby providing evidence that the PIP1 level is subject to proteasome-dependent ERAD. Given the established role of BiP chaperones in ER quality control and their functional coupling with ERAD, a ∼2-fold upregulation of *BiP1/2* transcripts (Figure S6) suggests an adaptive response to maintain ER homeostasis which is perturbed by an accumulation of misfolded or unassembled PIP1 proteins.

The lack of increased PIP1 accumulation in the *pip2;1 pip2;2 pip2;7 atg7* mutant may suggest that ERAD remains fully functional in this background and is sufficient to mediate PIP1 turnover. This observation is consistent with the established role of ERAD as a primary pathway for the degradation of misfolded or unassembled membrane proteins in plants (Duan et al, 2023; Wu et al, 2026). In contrast, autophagy is generally activated under severe or prolonged stress conditions, including abiotic and biotic challenges, where it contributes to bulk degradation and cellular homeostasis (Petersen et al, 2024). Together, these findings indicate that, under non-extreme conditions, ERAD alone may efficiently regulate PIP1 protein levels, while autophagy likely plays a supportive role during heightened stress. Top of Form

We refrained from generating ERAD- and autophagy-deficient double mutants due to concerns over plant viability and potential UPR activation, which could obscure PIP1 regulation.

### Physiological implication of PIP2-dependent regulation of PIP1s

The genetic loss of major PIP2 isoforms created an artificial situation which contrasts the high evolutionary constraints on the co-existence of plant PIP1 and PIP2 protein subclades. However, it may provide a blueprint to situations where *PIP2* expression is downregulated (Alexanderson et al, 2010; Hachez et al, 2014a; Chen et al, 2021). In such a scenario, the discovered function of PIP2s as obligatory, positive regulators of PIP1 expression may lead to degradation of newly synthesized PIP1 proteins and thereby coordinate PIP1 and PIP2 interaction and abundance. This regulation may provide at least two advantages, (i) a rapid control mechanism for water or other PIP-dependent membrane permeability, e.g., upon stress responses aiming at a rapid downregulation of PIPs, and (ii), a coordination of physiological properties that rely on and are established by PIP1-PIP2 heteromers (Bienert et al, 2018). Accordingly, already documented features of in planta PIP2 functions may also involve the contribution or co-regulation of PIP1 members: (i) under drought stress PIP2;1, PIP2;2, and PIP2;7 have been shown to be degraded already at the ER (Lee et al, 2009; Hachez et al, 2014a; Chen et al, 2021); similar results have been reported for other plants (Li et al, 2020; Chen et al, 2022). (ii) PIP2s have been associated with greater functional diversity in plants due to their flexible molecular regulation and broader substrate permeability (Maurel et al, 2021; Tyerman et al, 2021). (iii) Last not least, physiological and developmental roles of PIP2 isoforms in water relocation, lateral root development, or H_2_O_2_ signaling have been established using *pip2* loss-of-function mutants (Javot et al, 2003; Da Ines et al, 2010; Péret et al, 2012; Prado et al, 2013; Poitout et al, 2026). Vice versa, the discovery of PIP1-specific functions (Matsumoto et al, 2009; Postaire et al, 2010; Tian et al, 2016; Schley et al, 2025) may require considering the involvement and contribution of co-expressed PIP2 isoforms.

In summary, our findings indicate that PIP1 expression in Arabidopsis is dependent on PIP2s in an obligatory manner, and this regulation primarily involves the post-translational level. PIP2s, beyond their own aquaporin function, serve as regulatory scaffold for PIP1 stability and principal PIP1 expression. This adds an additional layer of regulatory interaction beyond the dependence of PIP1 isoforms on assistance by PIP2s to exit the ER by providing ER export motifs and/or masking PIP1 ER retention signals (Zelazny et al, 2009; Sorieul et al, 2011; Chevalier et al, 2014; Wang et al, 2020). The evolutionary conservation of the PIP1 and PIP2 subclades reflects this tight linkage of PIP1 and PIP2 isoform expression, although the specific implications and physiological contribution of PIP1 and/or PIP2 are still to be resolved (Mintseris and Weng, 2005; Soto et al, 2012; Bienert et al, 2018). These perspectives on PIP1–PIP2 interaction underscore the importance of considering the intrinsic PIP2-dependent control of PIP1 expression when studying PIP functions. Notably, genetic studies of PIP2 functions cannot be performed in isolation, since they are coupled to the concomitant regulation of PIP1 abundance. Although this intrinsic link has been analyzed here only for the model plant *A. thaliana*, we propose that it is likely observed also in other plants based on the persistence and conservation of PIP1 and PIP2 sequences (Soto et al, 2012; Bienert et al, 2018).

## Materials and methods

### Plant materials and growth conditions

Mutant lines were obtained from the Arabidopsis Stock Center (Sundaresan et al, 1995; Tissier et al, 1999; Scholl et al, 2000; Alonso et al, 2003) unless otherwise indicated. Wild-type and mutant *Arabidopsis thaliana* lines used in this study were of the Columbia (Col) ecotype, except for the *pip2;7-1* allele (Landsberg *erecta* (L*er*)) (Table S1). *hrd1a hrd1b bri1-9* was obtained from Linchuan Liu, Shanghai Plant Center for Stress Biology, CAS; Su et al, 2011); the loss of the *bri1-9* allele - after crosses to introgress *hrd1a hrd1b* - was assessed by a dCAPS marker (Jin et al, 2007; Table S3). Multiple mutant alleles were combined by genetic crossing or by inducing CRISPR/Cas9- dependent deletions; some multiple mutants had been described before (Schley et al, 2025; Poitout et al, 2026; Table S1).

*pip2;1 pip2;2 pip2;7-2 dln1-1* was generated by a CRISPR/Cas9-induced deletion in *DLN1*. Two gRNAs (Labun et al, 2019; Table S4) had been assembled (each twice) into the binary vector pDGE347 (Ordon et al, 2020; 2021) for *Agrobacterium*-mediated transformation (Clough and Bent, 1998). Homozygous T2 plants with *DLN1* deletions were confirmed by PCR and sequencing. *dln1- 1* had a 210 bp deletion (nucleotides 687–896 relative to the ATG start codon).

EGFP-tagged PIP1;2 and HA-tagged PIP1;1 or PIP1;2 under the control of their respective *PIP1* promoter were compiled into the binary vector pPm42GW3 (Karimi et al, 2005) (Table S4) and transformed into the respective *pip1;1* or *pip1;2* mutant. One homozygous, single insertion transgenic line harboring the EGFP- or the HA-tagged *PIP* version in *pip1;1* or *pip1;2* was crossed with the *pip1;1 pip2;1 pip2;2* or the *pip1;2 pip2;1 pip2;2*, respectively. Homozygous lines representing a pseudo wild-type situation and a *pip2;1 pip2;2* mutant background harboring the same tagged PIP1 transgene were selected from the segregating populations. Thus, the expression of the tagged PIP1;1 or PIP1;2 proteins could be compared in these different backgrounds. The *pip1;2-2 pip2;1-2 pip2;2-3 pip2;7-5* (EGFP-PIP1;2) line was created by CRISPR/ Cas9-induced deletions of *PIP2;7* in the same manner as described for *DLN1* deletions. *pip2;7-5* showed a 272 bp deletion (nucleotides 155 to 426 were deleted with A of the start codon being nucleotide 1) in the first and second exon and introduced a frameshift.

Plants were grown on half-strength Murashige and Skoog (MS) agar plates with a 16 h light (100- 120 μE m^-2^ s^-1^ fluorescent light) / 8 h darkness regime at 22°C and 60% relative humidity for 14 days to obtain rosette tissue for microsomal membrane fractions isolation and for gene expression analysis. Plants were grown on soil with a 10 h light at 100-120 μE m^-2^ s^-1^ /14 h darkness regime at 22°C and 60% relative humidity for RT-qPCR analysis of *PIP1* gene expression (21-day-old), for polysome profiling (21-day-old) and co-immunoprecipitation (28-day-old), for protoplast isolation (4- 5-week-old), or for stable transformation.

### Cloning

BiFC constructs were cloned according to Grefen and Blatt (2012) (Figure S9; Table S2; Table S4). Plasmids containing an N-terminal EGFP- or HA-tag to *PIP1;1* and *PIP1;2* coding sequences (with a linker encoding glycine-alanine in front of the PIP1 ATG start codon) under the control of the respective *PIP1* promoters and 3’-UTRs that were generated by PCR-based joining of fragments and a GATEWAY^TM^ two-fragment vector recombination method (Karimi et al, 2005; Karimi et al, 2007). pDONRP4P1R and pDONR221 containing different fragments were then recombined into pPm42GW3-*At2s3_pro_*::GFP for stable transformation (Bensmihen et al, 2004; Figure S10; Table S3). *35S_pro_::PIP1;2-EGFP* was established in pDONRP4P1R and used for transient expression in protoplasts (Figure S11; Table S4). *PIP2;1* full-length cDNA was amplified by PCR, inserted into pDONR221, and recombined into pBS-2x*35S*-HA-GW-tNOS (Table S2, S4).

### Microsomal fractions

Microsomal fractions were isolated from about 0.5 g rosette leaves homogenized in 30 ml 50 mM HEPES-KOH, pH 7.5, 5 mM EDTA, 0.5 M sucrose, 2 mM DTT, protease inhibitors, 0.1% PVPP, 0.1 mg ml ^-1^ BHT using a mortar and pestle. After centrifugation at 8000 × g for 10 min at 4°C, the supernatant was filtered through Miracloth^TM^ and centrifuged at 110000 × g for 1 h at 4°C. The pellet was resuspended in 100 µl of 0.33 M sucrose, 5 mM K₃PO₄, 4 mM KCl, 2 mM DTT, homogenized in a douncer and stored at −80°C in a final volume of ∼300 µl.

### Antibodies, ELISA, and Western blotting

Rabbit anti-PIP1 antiserum (Henzler et al, 1999; Figure S2), rabbit anti-PIP2-antiserum raised against C-terminal peptide RASGSKSLGSFRSAANV coupled to keyhole limpet hemocyanin detecting PIP2;1, PIP2;2, PIP2;3 (Da Ines, 2008; the peptide used to raise antibodies is identical in all three isoforms, however, PIP2;3 is very weakly expressed: thus, the lack of a signal in the *pip2;1 pip2;2* mutant background (Fig. 3A, Fig. S5) does not necessarily exclude the presence of PIP2;3), mouse anti-HA antibody (H3663, Sigma, Germany), anti-mouse cy3-linked and anti-rabbit cy5-linked antibodies (Amersham Biosciences, Germany) were used for immunological detection.

ELISA was performed according to Santoni et al (2006) with modifications. The samples (in triplicate) were serially diluted with sodium carbonate buffer (pH 9.5) using 96-well plates (NUNC Immuno Plate Maxisorb). The 42 N-terminal amino acids of PIP1;1 served as a peptide standard. Rabbit anti- PIP1 antiserum and horseradish peroxidase-conjugated anti-rabbit-IgG using 2,2′-azino-bis (3- ethylbenzothiazoline-6-sulfonic acid) as substrate were used for quantification. Background binding was determined using primary anti-PIP1 antiserum blocked with the antigenic 42 amino acid-long PIP1;1 peptide. Data were analyzed with a 5-parameter logistic regression (http://elisaanalysis.com//app).

For Western blotting, 1.5 µg microsomal protein were separated on 10% SDS-polyacrylamide gels. PIP1 isoforms were detected using the anti-PIP1 antiserum (Henzler et al, 1999; Fig. S2). Quantification was performed using ImageJ (Schindelin et al, 2012).

### Mass spectrometry

Microsomal fractions were prepared from rosette leaves of 28-day-old plants grown on soil and from roots harvested from plants grown in half-strength MS medium shaken for 21 days in the dark. Proteins were denatured in Laemmli buffer with 100 mM DTT added. Pull-down samples and microsomal fractions were digested using a modified FASP protocol (Wiśniewski et al, 2009; Grosche et al, 2015). Proteins were reduced and alkylated with DTT and iodoacetamide and filtered through a 30 kDa cut-off filter device (PALL). After sequential washes (8 M urea, 0.1 M Tris-HCl pH 8.5, 50 mM ammonium bicarbonate), digestion was performed on-filter using 1 µg Lys-C (Wako Chemicals) (2 h, 22°C) and 2 µg trypsin (Promega) (16 h, 37°C). Peptides were collected by centrifugation (10 min, 14000 × g), acidified (0.5% trifluoroacetic acid), and stored at −20°C. Following centrifugation (5 min, 4°C), LC-MS/MS analysis was conducted (Hauck et al, 2010) using a nano-HPLC (Dionex) coupled to an LTQ-OrbitrapXL mass spectrometer (Thermo Scientific). Approximately 0.5 µg of each sample (5% acetonitrile (ACN), 0.1% formic acid) was injected and loaded onto a trap column (100 µm inner diameter, 2 cm long, Acclaim PepMap100 C18, 5 µm, 100 Å) at 30 µl/min. After 5 min, peptides were eluted and separated on an analytical column (75 µm inner diameter × 15 cm, Acclaim PepMap100 C18, 3 µm, 100 Å) using a 140 min gradient (5–30% ACN), followed by a 5 min gradient (30–95% ACN). Between each sample, the gradient was set back to 5% ACN and left to equilibrate for 20 minutes. The 10 most abundant peptide ions of the MS prescan were selected for fragmentation in the linear ion trap if they were at least doubly charged. During fragment analysis a high-resolution (60,000 full width at half maximum) MS spectrum was acquired over a mass range of 300–1500 Da.

### Label-free analysis using Progenesis LC-MS

The acquired spectra were loaded to the Progenesis QI software (version 2.0, Nonlinear Dynamics) for label free quantification and analyzed as described previously (Hauck et al, 2010; Merl et al, 2012; Molin et al, 2014). Retention times were aligned by automatic alignment to a maximal overlay of all features. MS/MS spectra of features with charges of +2 to +7 were exported as Mascot generic file and used for peptide identification with Mascot (version 2.5) in the TAIR protein database (version 10, 35386 sequences, 14483180 residues). Search parameters used were: 10 ppm peptide mass tolerance and 0.6 Da fragment mass tolerance, one missed cleavage allowed, carbamidomethylation was set as fixed modification, methionine oxidation and asparagine or glutamine deamidation were allowed as variable modifications. Searches were filtered with a Mascot ion score of 30 and an appropriate significance threshold p to reach a maximum false discovery rate of 1%. Peptide assignments were re-imported into the Progenesis QI software and the spectral counts of all peptides allocated to each protein were summed up per sample. Normalized proteins abundances and spectral counts were exported and used.

### Protoplast isolation and PEG-mediated transformation

Protoplasts were isolated from leaves from 4- to 5-week-old short-day-grown plants (Wu et al (2009). The adaxial epidermis was peeled, and leaves were digested in 10 ml enzyme solution. After 60-90 min, protoplasts were released, collected, centrifuged at 100 g for 2 min, and washed. Two × 10⁵ cells ml ^-1^ were used for transformation (Yoo et al, 2007).

### Reverse transcription and quantitative PCR

RNA was extracted from rosette leaves using the innuPREP RNA Kit (Analytik Jena, Germany). One µg RNA was treated with DNase I and reverse transcribed for cDNA synthesis using QuantiTect Reverse Transcription Kit (Qiagen, Netherlands). Gene expression data obtained by real-time PCR quantification (Applied Biosystems 7500) were normalized to stably expressed transcripts *UBIQUITIN5* and *S16* (Vandesompele et al, 2002) (Table S3).

### Polysome profiling

The protocol followed in this study is adapted from Mustroph et al (2009) and Lecampion et al (2016). Twenty-one-day-old rosette leaves were flash-frozen in liquid N₂, and ground into powder. Approximately 3.6 g of tissue was homogenized in polysome extraction buffer and centrifuged at 16,000 × g for 15 min at 4°C. The supernatant was purified using a sucrose cushion centrifuged at 230000 × g for 3 h, yielding a ribosome-rich pellet. Polysomes were obtained by loading 500 µl resuspended ribosomal pellet onto a 20–50% sucrose gradient and centrifuging at 182,000 × g for 2.5 h at 4 °C. Fractions (1.5 ml) were collected and RNA concentration recorded (OD at 260 nm). For mRNA extraction, 4 µl 1% linear polyacrylamide, 8 M guanidine hydrochloride, and ethanol were added to the polysome fraction, followed by overnight precipitation at −20 °C. After centrifugation (47,800 × g, 45 min) RNA was purified using the innuPREP RNA Kit (Analytik Jena, Jena, Germany). Two µg RNA were treated with DNase I and used for cDNA synthesis (Invitrogen). *PIP1* gene expression was assessed by real-time PCR quantification (Applied Biosystems 7500) and normalized using stably expressed genes *TUBULIN6* and *RHIP1* selected from a pool of references also including *UBIQUITIN5*, *GRF1*, *EF1ALPHA*, and *S16* (Vandesompele et al, 2002) (Table S5).

### Co-immunoprecipitation assay

Co-immunoprecipitation was conducted as described by Zelazny et al (2007) with minor modifications. Microsomal fractions (∼150 μg) were prepared and solubilized in 250 μl buffer (0.02 M Tris, 0.136 M NaCl, pH 7.6) containing 1%, 2%, and 3.5% n-octyl-β-D-thioglucopyranoside for 4 h at room temperature on a rotating wheel. After centrifugation (169,000 g, 40 min, 4°C), the supernatant was incubated with 25 μl protein A-agarose (Roche, Germany) for 1 h at 4°C to address nonspecific binding. After centrifugation (12,000 × g, 1 min, 4°C), the supernatant was incubated overnight at 4°C with 1 μl mouse anti-HA antibodies (H3663, Sigma, Germany). Protein A-agarose (50 μl; Roche, Germany) was added and incubated for 4 h at 4°C. The agarose-antibody-antigen complexes were collected by centrifugation (12,000 × g, 1 min, 4°C), washed four times with solubilization buffer and four times with TBS buffer. The resin was then incubated in 60 μl 1× Laemmli buffer with 100 mM DTT and 2% SDS at 70°C for 10 min. Proteins were resolved on 15% SDS polyacrylamide gels and transferred to a PVDF membrane. Western blot analyses used anti-HA (H3663, Sigma, Germany) or rabbit anti-PIP2;1/PIP2;2/PIP2;3 antiserum (see above).

### Pharmacological inhibition experiments

For MG132 treatment, protoplasts were isolated from stable transgenic *pip1;2 pip2;1 pip2;2 pip2;7* (EGFP-PIP1;2) lines and maintained at 2 × 10⁵ cells/ ml in W5 solution. Ninety µl of protoplast solution was mixed with 10 µl of 500 µM MG132 (final 50 µM). Controls received 10 µl DMSO. Samples were incubated horizontally in the dark for 18 h before confocal laser scanning microscopy (SP8, Leica, Germany). EGFP-PIP1;2 fluorescence was measured (488/500-550 nm excitation/emission, 2% laser intensity, 20X objective). Z-stack images (3 µm intervals) were processed using ImageJ for maximum intensity projections.

## Acknowledgements

We greatly acknowledge Olivier Da Ines who originally discovered the phenomenon of PIP1 downregulation when studying the *pip2;1 pip2;2* mutant. KJ acknowledges a doctoral fellowship of the German DAAD. CL was supported by a doctoral fellowship of the China Scholarship Council (CSC). Valentine Haury, Franziska Bottler and Lea Korn were involved in several steps of characterization of the *pip* and *dln* mutants. We thank the teams of Roland Beckmann (Gene Center, LMU, Munich) and Mark Stitt (Max Planck Institute, Potsdam) for guiding us with the polysome profiling experiment.

## Author contributions

KJ, JL, CL, and ARS planned and designed the research. KJ, CL, JL, BG, and ARS performed the experiments. JZ and JMP contributed proteomic analyses. KJ, CL, and ARS wrote the article.

## Supplementary material

Supplementary material is available online.

**Figure S1**. Images of 14-day-old rosette of WT, *pip2;1 pip2;2*, *pip2;1 pip2;2 pip2;7*, *pip2;1 pip2;2 pip2;4 pip2;6*, *pip2;1 pip2;2 pip2;4 pip2;6 pip2;7* grown on soil.

**Figure S2**. PIP1 isoform detection by rabbit anti-PIP1 antiserum.

**Figure S3**. CBB stained SDS gels used for normalization of Western blots.

**Figure S4.** The representative images of the protoplasts expressing PIP1;2-EGFP alone and co- expression of PIP1;2-EGFP and HA-PIP2;1 used for quantification of PIP1;2-EGFP fluorescence.

**Figure S5.** PIP1 and PIP2 proteins interact with each other impacting the targeting of PIP1.

**Figure S6.** *UPR* gene expression analysis of *pip2;1 pip2;2 pip2;7* mutants by RT-qPCR.

**Figure S7.** ELISA for ERAD-defective mutants reveals a partial role of ERAD.

**Figure S8.** ERAD is partially involved in repressing PIP1 protein levels in absence of PIP2;1, PIP2;2, and PIP2;7.

**Figure S9.** Cloning strategy of BiFC expression constructs.

**Figure S10.** Cloning strategy of N-terminal EGFP/HA-tagged PIP1;1/ PIP1;2 constructs.

**Figure S11.** Cloning strategy of C-terminal EGFP-tagged PIP1;2 construct.

**Table S1.** Mutant and transgenic lines.

**Table S2.** Plasmids.

**Table S3.** Oligonucleotides used for genotyping.

**Table S4.** Oligonucleotides used for cloning.

**Table S5.** Oligonucleotides used for RT-qPCR.

## Conflict of interest

All authors declare no competing interests.

## Data availability

All relevant data can be found within the article and its supporting materials. Materials will be available through the corresponding authors and Arabidopsis stock centers.

## Notes

### Competing Interest Statement

The authors have declared no competing interest.

