## Supplementary Figures & Tables for "Concomitant post-translational repression of *Arabidopsis* PIP1 aquaporins upon the loss of major PIP2 isoforms"

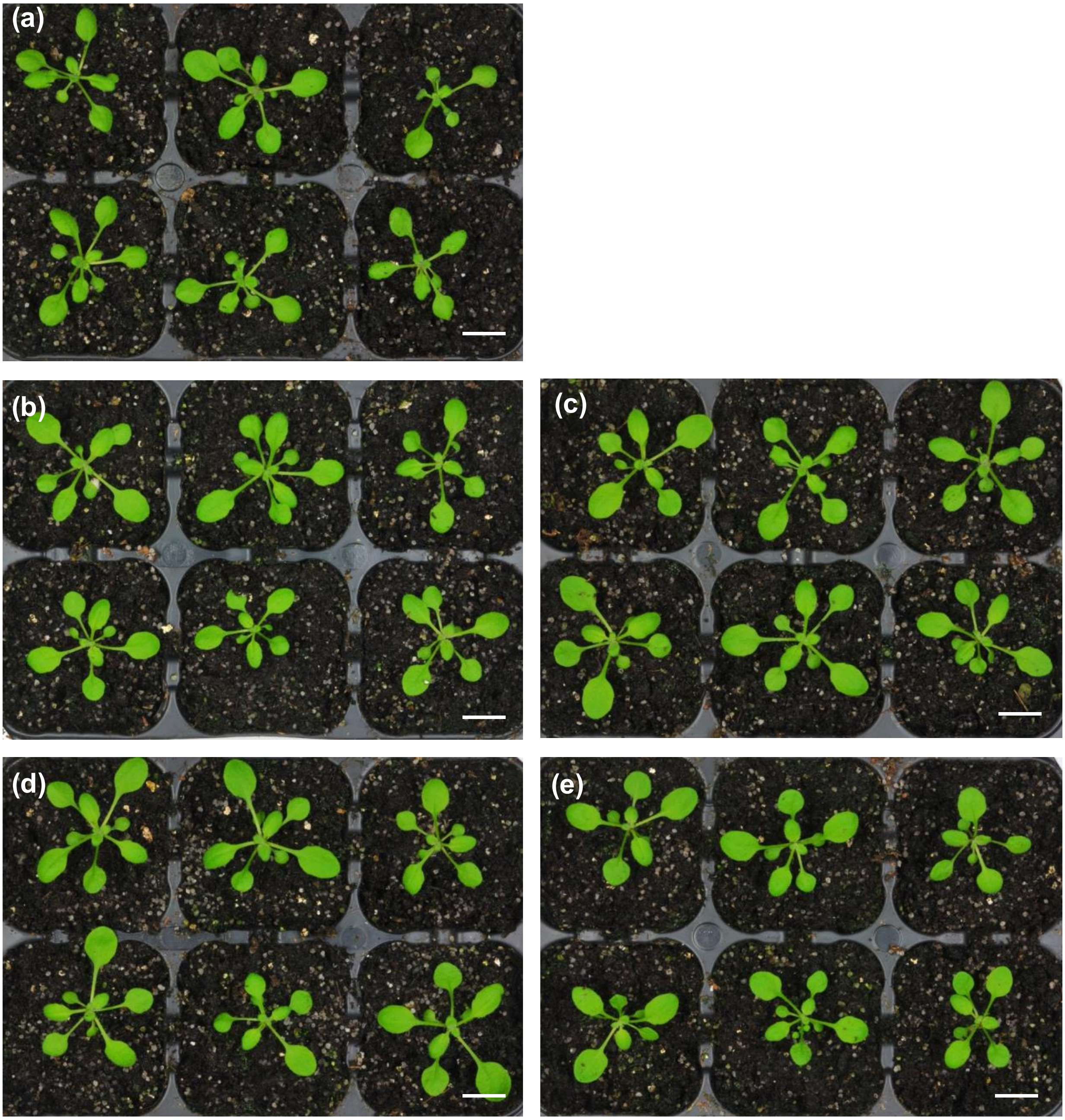

**Figure S1**. Images of 14-day-old rosettes of (a) WT, (b) *pip2;1 pip2;2,* (c) *pip2;1 pip2;2 pip2;7,* (d) *pip2;1 pip2;2 pip2;4 pip2;6,* and (e) *pip2;1 pip2;2 pip2;4 pip2;6 pip2;7* grown on soil (20-22 °C, 16 h fluorescent light, 80% relative humidity). Bars equal 1 cm.

(a)

```
AtPIP1;1_At3g61430  MEGKEEDVRVGANKFPERQPIGTSAQS-DKDYKEPPPAPFFEP
AtPIP1;2_At2g45960  MEGKEEDVRVGANKFPERQPIGTSAQS-DKDYKEPPPAPLFEP
AtPIP1;3_At1g01620  MEGKEEDVRVGANKFPERQPIGTSAQT-DKDYKEPPPAPFFEP
AtPIP1;4_At4g00430  MEGKEEDVRVGANKFPERQPIGTSAQSTDKDYKEPPPAPLFEP
AtPIP1;5_At4g23400  MEGKEEDVNVGANKFPERQPIGTAAQTESKDYKEPPPAPFFEP
```

(b)

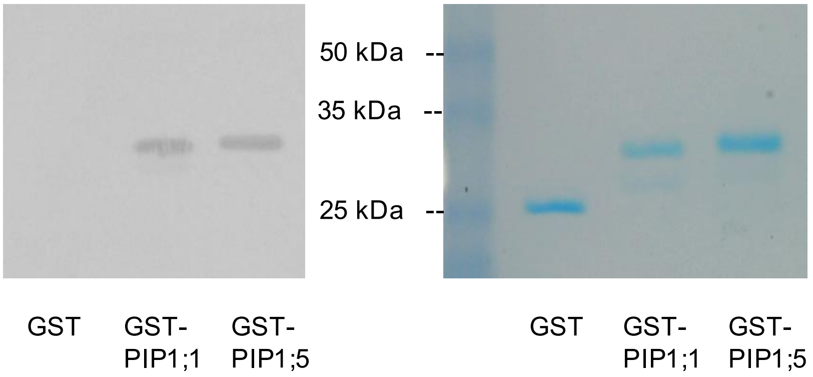

**Figure S2**. PIP1 isoform detection by rabbit anti-PIP1 antiserum. Rabbit anti-PIP1 antiserum was raised using recombinant GST-PIP1;1 fusing the N-terminal 42 amino acids of PIP1;1 to GST (Henzler et al., 1999). This region is highly homologous among all other PIP1 isoforms, however, PIP1;5 harbors two deviating subregions. Therefore, the polyclonal anti-PIP1 antiserum was tested against GST-PIP1;5 (encompassing 44 N-terminal residues) in comparison to GST and GST-PIP1;1. The detection of GST-PIP1;5 indicated that all five PIP1 isoforms are detected by the antiserum.

(a) Alignment of the N-termini of all *A. thaliana* PIP1 isoforms. The PIP1;1 epitope used to raise antibodies is underlined. Amino acid residues identical to PIP1;1 are indicated in blue, divergent residues (including conserved S/T changes) are highlighted in red.

(b) Proteins were electrophoresed using a 12% SDS-polyacrylamide gel. Western blot was developed with a 1:10,000-diluted rabbit anti-PIP1 antiserum, which had been preincubated with 20 µg $ml^{-1}$ recombinant GST protein to block reaction with the GST entity of the fusion proteins (note that the carrier protein GST was no more detected by the Western blot analysis). Anti-rabbit horseradish peroxidase-coupled antibodies and DAB staining was used for detection. A parallel SDS polyacrylamide gel was stained with Coomassie Brilliant Blue G250 to indicate the protein loading.

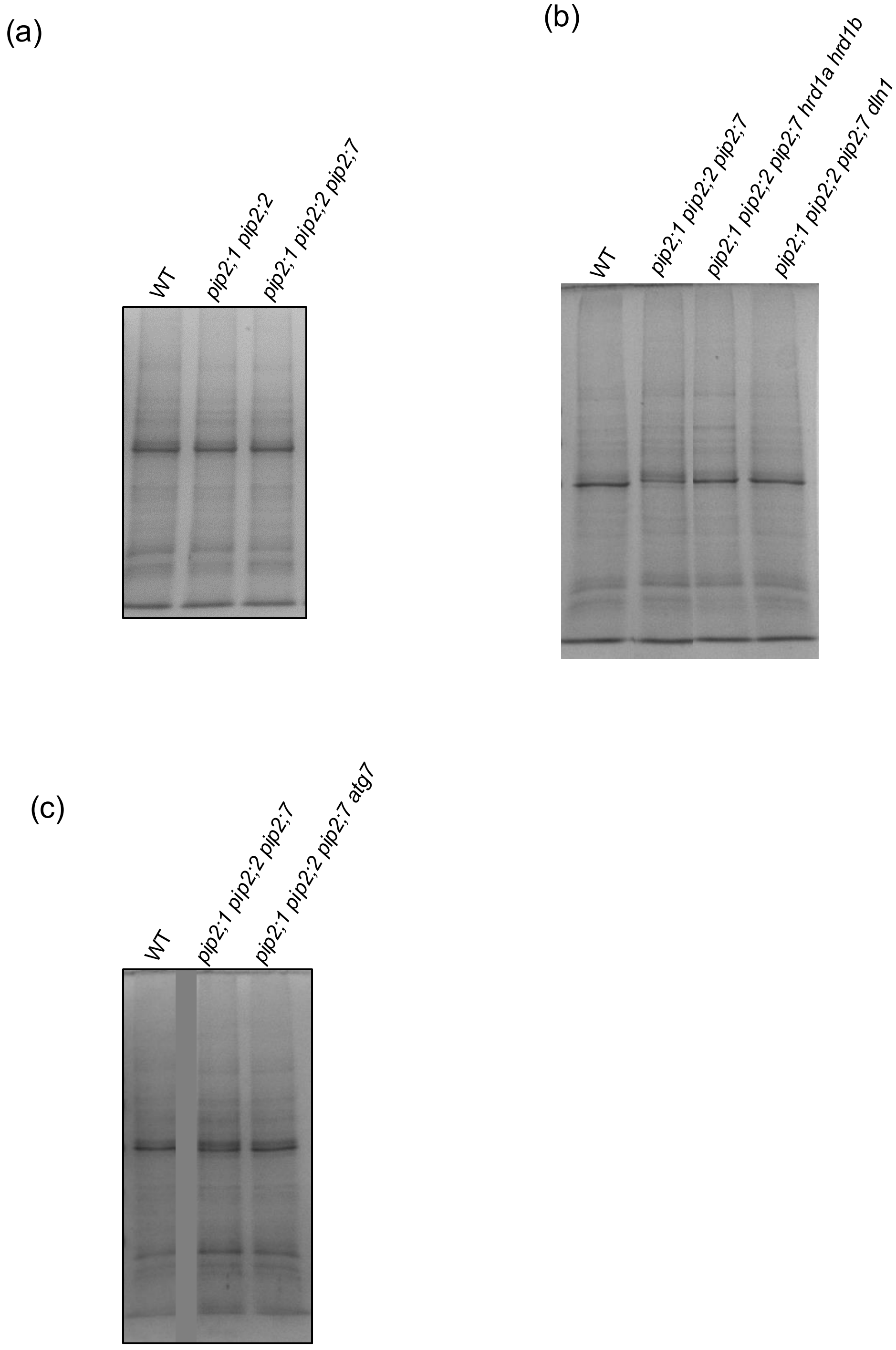

**Figure S3**. Representative Coomassie-stained SDS polyacrylamide gels used for normalization of Western blots shown in (a) Fig. 1b, (b) Fig. 5b, and (c) Fig. 6a.

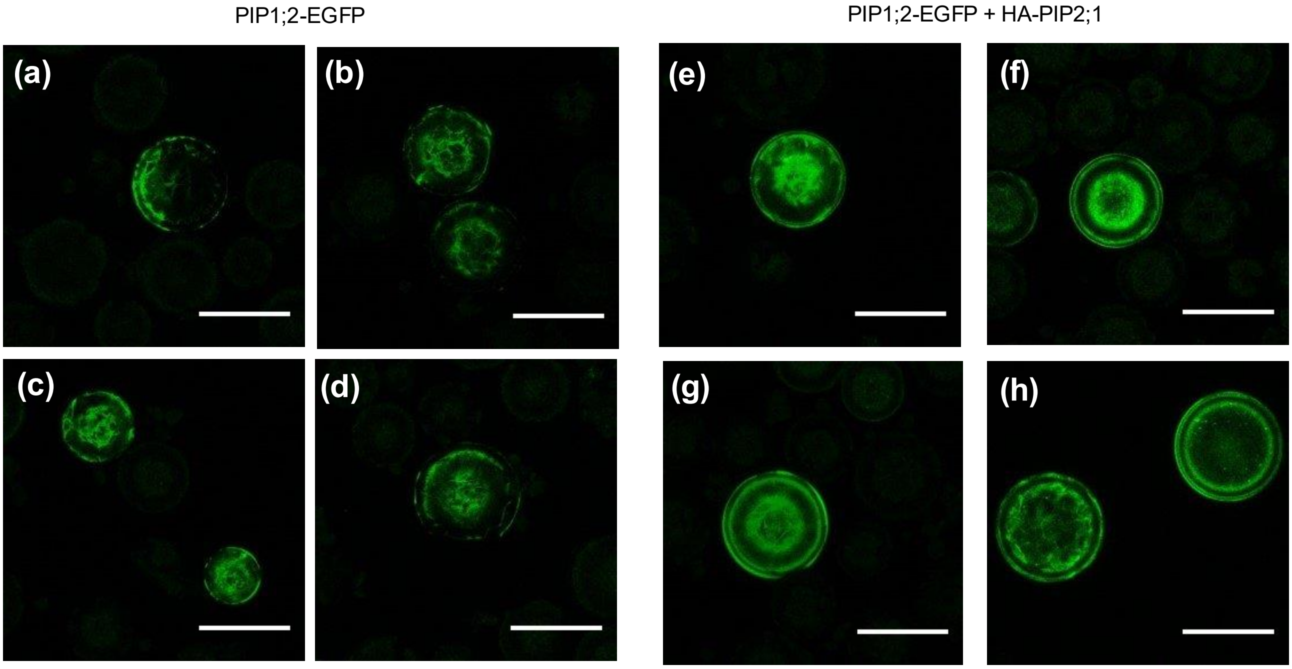

**Figure S4.** Representative images of protoplasts expressing PIP1;2-EGFP alone (a, b, c, d) and co-expression of PIP1;2-EGFP and HA-PIP2;1 (e, f, g, h). Such images were used for quantification of PIP1;2-EGFP fluorescence shown in Fig. 2c. Twenty-two images of the protoplasts were taken at a Z interval of 5 µm. Image J software (Fiji) was used to create sum of slice projections of the Z-stack images of the protoplasts. Scale bars, 50 µm.

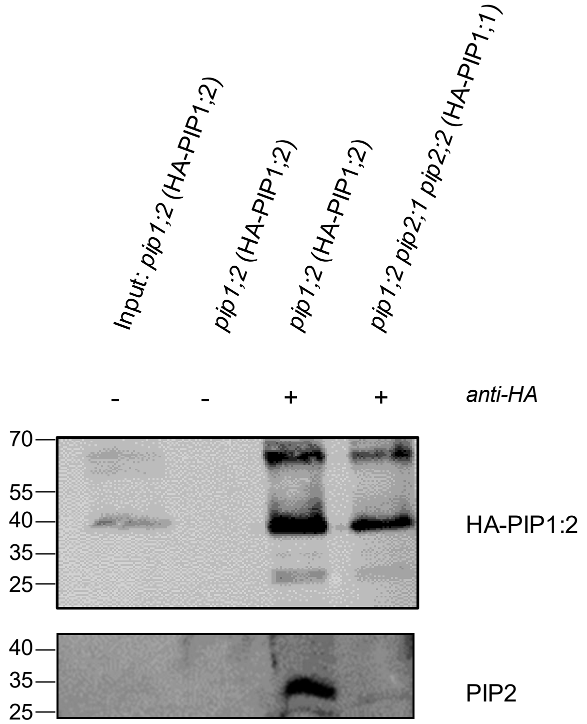

**Figure S5.** PIP1 and PIP2 proteins interact with each other impacting the targeting of PIP1.

Microsomal membrane fractions extracted from 28-day-old rosette leaves of transgenic *pip1;2* (*PIP1;2$_{pro}$::HA-PIP1;2*) and *pip1;2 pip2;1 pip2;2* (*PIP1;2$_{pro}$::HA-PIP1;2*) were subjected to immunoprecipitation using an anti-HA antibody and analyzed by Western blotting using the anti-HA antibody (upper panel, detecting HA-PIP1;2 monomer and dimer at about 36 kDa and 65 kDa) or an anti-PIP2;1/PIP2;2/PIP2;3 antiserum (lower panel, detecting 28 kDa PIP2 monomers). +, anti-HA antibody was added for immunoprecipitation; – antibody was absent during immunoprecipitation.

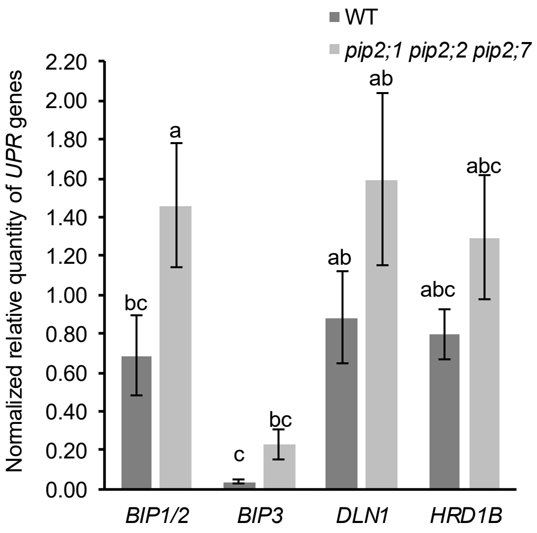

**Figure S6**. *UPR* gene expression analysis of *pip2;1 pip2;2 pip2;7* mutants by RT-qPCR. RNA was isolated from 14-day-old rosette tissue of WT and *pip2;1 pip2;2 pip2;7* grown on one half strength Murashige and Skoog with a 16-h-light/8-h-dark cycle mutant. RT-qPCR was performed. The sequences of *BIP1 and BiP2* genes are nearly identical, therefore a single primer pair detecting both genes had been employed. Data are represented after normalization to reference genes. Bars are means ± SD of three independent experiments. One-way ANOVA with Tukey's post hoc method was used to examine significant differences (P< 0.050) in pairwise comparison and classified by letters. WT, wild type.

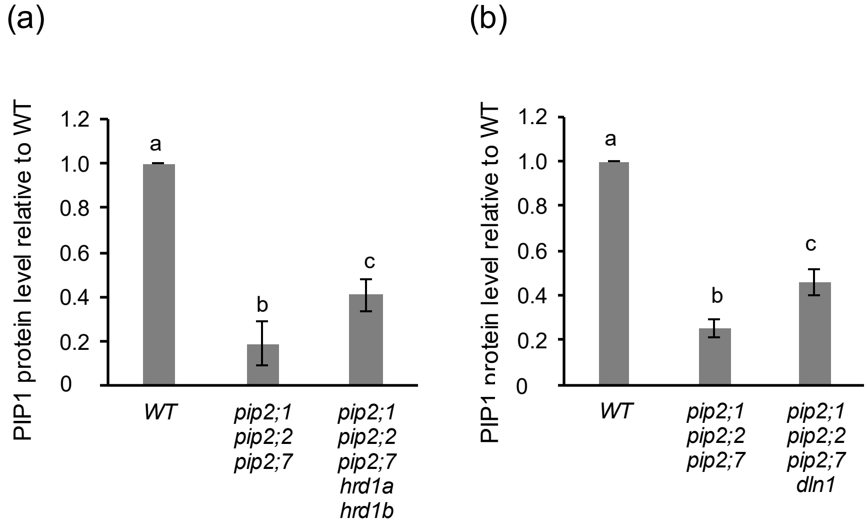

**Figure S7**. ELISA for ERAD defective mutants reveals partial role of ERAD.
Microsomal fraction protein was isolated from 14-day-old rosette tissue of plants grown on agar plates. Anti-PIP1 antiserum detecting all five PIP1 isoforms (Fig. S2) was used to probe PIP1.
(a) Introgression of *hrd1a hrd1b* knockout.
(b) Introgression of *dln1* knockout.
Bars are means ± SD of three independent experiments each. One-way ANOVA with Tukey's post hoc method was used to examine significant differences (P< 0.050) in pairwise comparison and classified by letters. WT, wild type.

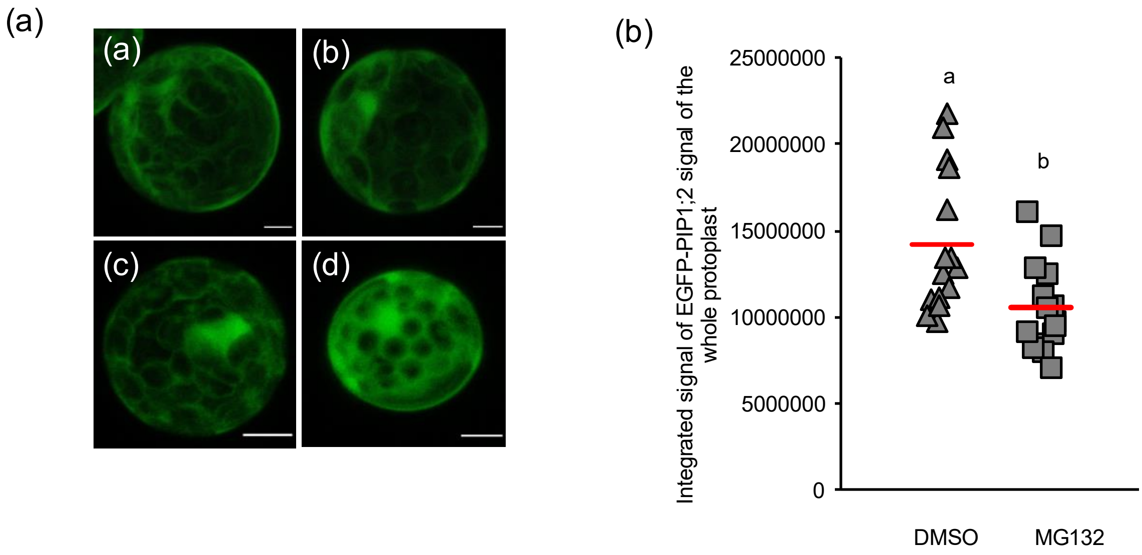

**Figure S8**. ERAD is partially involved in repressing PIP1 protein levels in absence of PIP2;1, PIP2;2, and PIP2;7.
(a) Additional representative images of protoplasts (in addition to Figure 5d) showing an enhanced cytoplasmic accumulation of EGFP-PIP1;2 signal upon MG132 treatment (c, d) compared to DMSO control (a, b). Scale bars, 50 µm.
(b) Quantification of integrated EGFP-PIP1;2 fluorescence intensities from whole protoplasts as shown in Figure 5d and Figure S8a. Data are plotted as integrated density of the EGFP-PIP1;2 signal from protoplasts treated with DMSO and MG132 (n = 15 protoplasts). Data are represented as scatter plot with the red horizontal line defining the mean value. Mann-Whitney Rank sum test was performed to examine significant differences ($P < 0.050$) classified by letters.

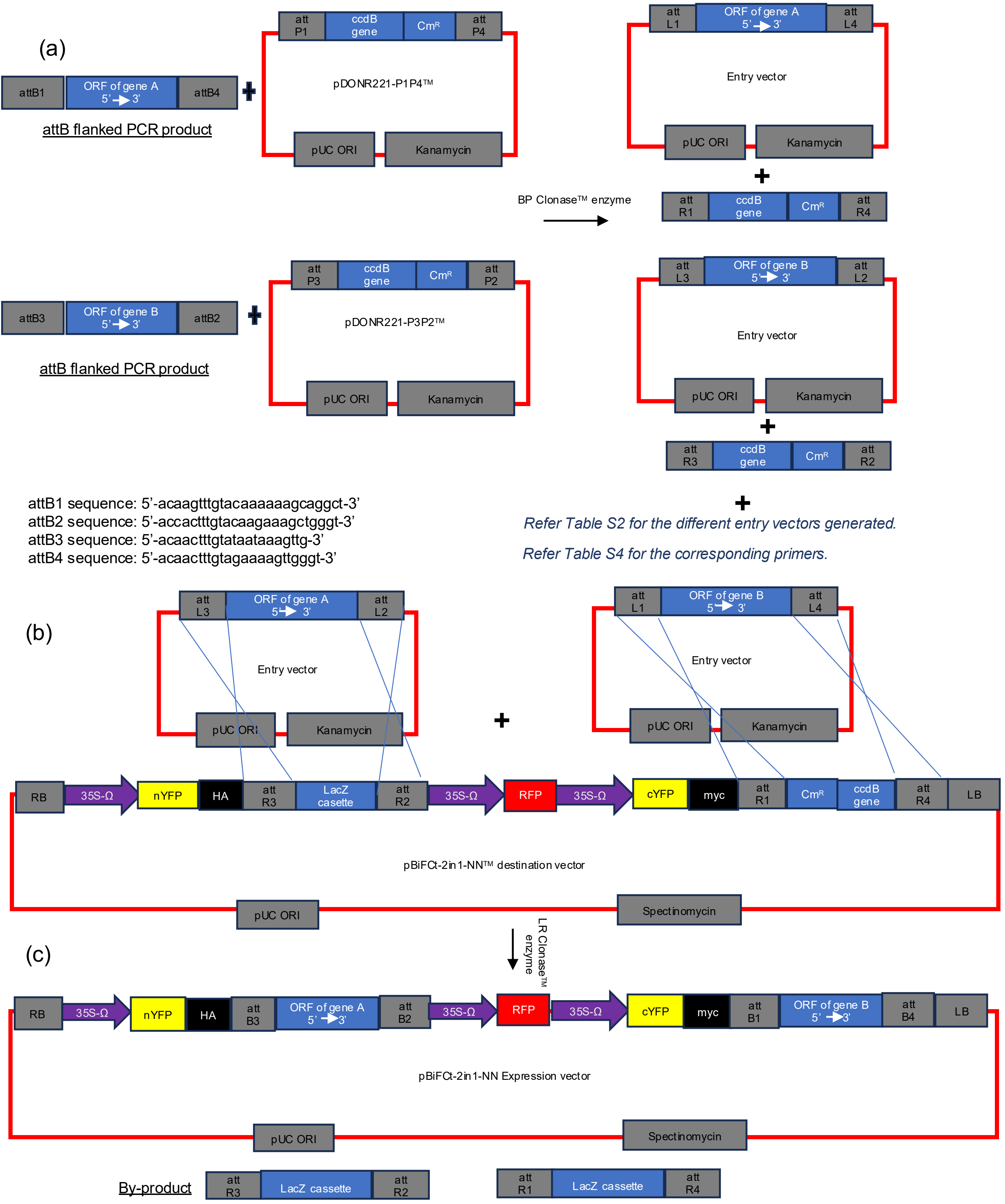

**Figure S9.** Schematic representation of the cloning strategy of BiFC expression constructs (Table S2). (a) BP recombination with the attB-flanked coding sequences (open-reading-frame, ORF) of gene A and gene B and the donor vectors pDONR221-P1P4 ™ and pDONR221-P3P2 ™ generates the entry vectors. (b) LR recombination of the entry vectors with the pBiFCt-2in1-NN destination vector (Blatt & Grefen, 2012) generates the expression vector with the genes A and B tagged with N-terminal YFP and C-terminal YFP fragments, respectively. ccd B gene: gyrase inhibitor gene; Cm$^R$ gene: chloramphenicol acetyl transferase gene; RB: T-DNA right border; LB: T-DNA left border.

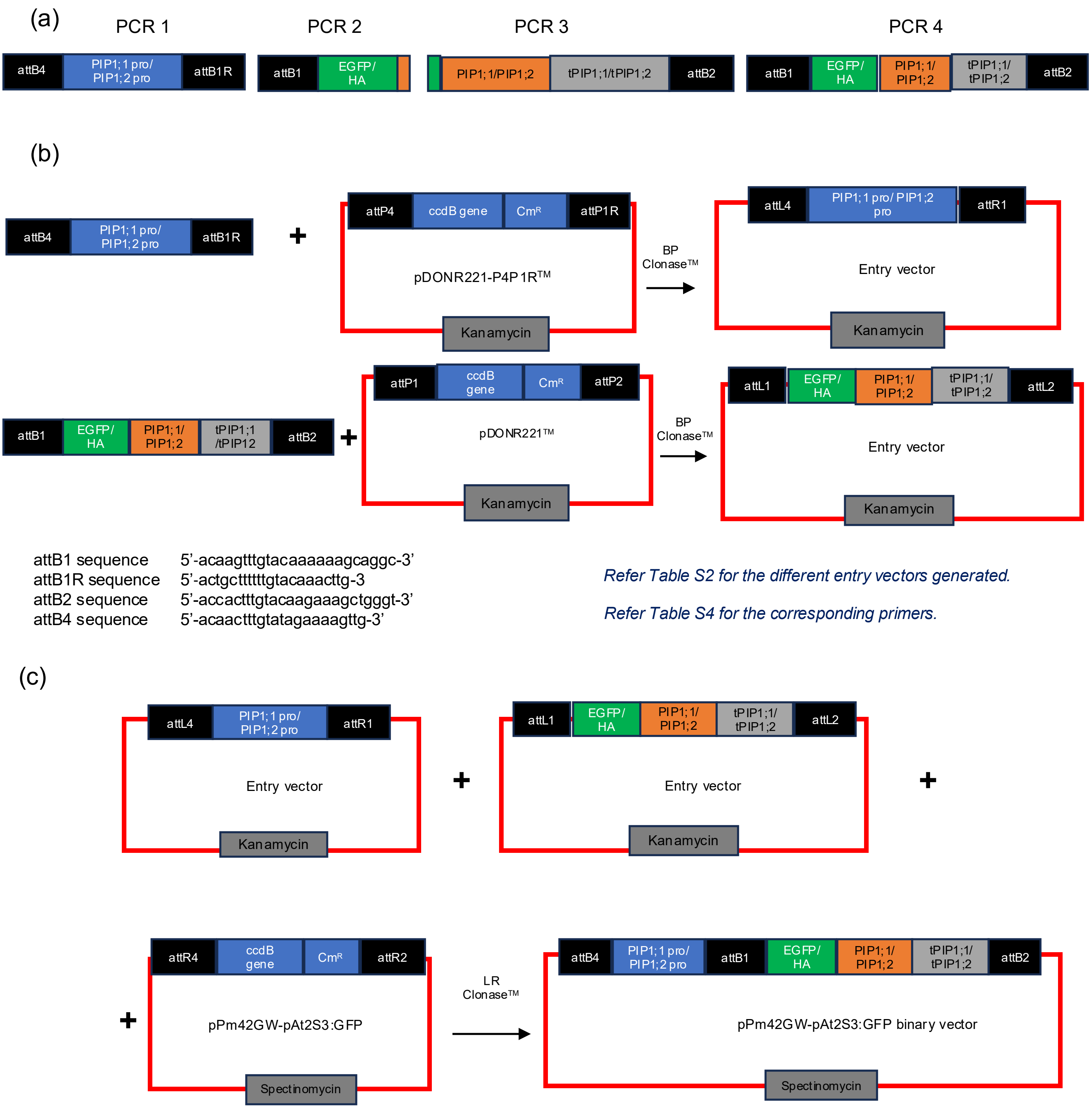

**Figure S10**. Schematic representation of the construction of N-terminal EGFP/HA-tagged PIP1;1/ PIP1;2 fusions (Karimi et al., 2005; Karimi et al., 2007).

(a) PCR-amplified fragments were created with attB primer extensions in PCR1 and PCR4. PCR4 was performed with overlapping PCR2 and PCR3 fragments as template and PCR2 forward and PCR3 reverse oligonucleotides (Table S2, S4).

(b) The attB-containing fragments were then assembled into the pDONRP4P1R and pDONR221 through BP recombination.

(c) Finally, for stable transformation these entry vectors were recombined into the binary pPm42GW-pAt2s3:GFP vector through LR recombination reaction.

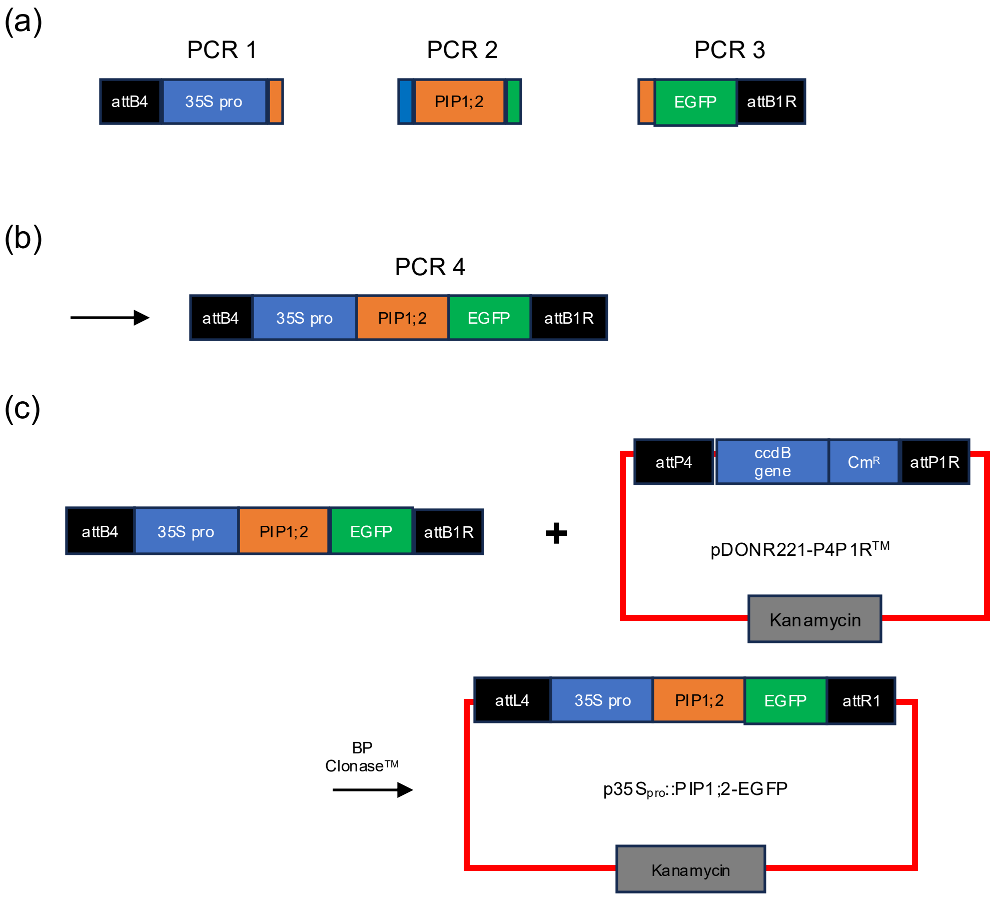

attB1R sequence 5'-actgctttttgtacaaacttg-3
attB4 sequence 5'-acaactttgtatagaaaagttg-3'

*Refer Table S2 for the different entry vectors generated.*

*Refer Table S4 for the corresponding primers.*

**Figure S11**. Schematic representation of the construction of the C-terminal EGFP-tagged PIP1;2 fusion used for transient expression (Karimi et al., 2005; Karimi et al., 2007).
(a) Modules of CaMV 35S$_{pro}$, PIP1;2 coding sequence, and EGFP were amplified with overlaps to the neighboring modules by PCR (PCR1, 2, and 3).
(b) These fragment were used as templates to create the fusion with attB4 and attB1R primer extensions by PCR4 (Table S2, S4).
(c) The attB-containing fragments were assembled into the pDONRP4P1R through BP recombination to create p35Spro::PIP1;2-EGFP.

**Table S1**. Mutant and transgenic lines.

| Allele(s) | Mutant /Transgenic line | Specification | Reference |
|---|---|---|---|
| *atg7-3* | *pip2;1-2 pip2;2-3 pip2;7-2 atg7-3* | Generated by crossing of *pip2;1-2 pip2;2-3 pip2;7-2* with *pip2;1-2 pip2;2-3 atg7-3* which was generated by crossing of *pip2;1-2 pip2;2-3* with *atg7-3* (SAIL_11H07, Thompson et al., 2005). | This work |
| *dln1-1* | *pip2;1-2 pip2;2-3 pip2;7-2 dln1-1* | CRISPR/Cas9 addressing *DLN1* in *pip2;1-2 pip2;2-3 pip2;7-2*. The generated deletion was confirmed by PCR and sequencing indicating a 210 bp deletion (nucleotides 687–896 relative to the ATG start codon). | This work |
| *hrd1a hrd1b* | *pip2;1-2 pip2;2-3 pip2;7-1 hrd1a hrd1b* | Generated by crossing of *pip2;1-2 pip2;2-3 pip2;7-1* (*pip2;7-1* allele is based on CSHL_GT19652 insertion line) with *pip2;1-2 pip2;2-3 hrd1a hrd1b* which had been generated by crossing *pip2;1-2 pip2;2-3* with *hrd1a hrd1b bri1-9* (this triple mutant based on SALK insertion lines had been obtained from Linchuan Liu, Shanghai Plant Center for Stress Biology, CAS) (Su *et al.,* 2011). *BRI1-9* wild-type alleles were screened by using a dCAPS primers designed to introduce a *DraI* restriction site in *bri1-9* but not in the wild-type DNA sequence. | This work |
| *pip1;1-1 pip1;2-2 pip2;1-2 pip2;2-3 pip2;4-1 pip2;6-3 pip2;7-1 pip2;7-2 pip2;7-4 pip2;7-5* | **1.** *pip1;1-1* | **1.** GABI_437B11 | **1.**Rosso et al., 2003 |
| | **2.** *pip1;2-2* | **2.** SALK_019794 | **2.** Alonso et al., 2003 |
| | **3.** *pip2;1-2 pip2;2-3* | **3.** *pip2;1-2* (SM_3_35928, Tissier et al., 1999) and *pip2;2-3* (SAIL_169A03, Alonso et al., 2003) were used for crossing. | **3.** Da Ines, 2008 |
| | **4.** *pip1;2-2 pip2;1-2 pip2;2-3* | **4.** *pip1;2-2* and *pip2;1-2 pip2;2-3* were used for crossing. | **4.** This work |
| | **5.** *pip2;1-2 pip2;2-3 pip2;7-1* | **5.** *pip2;7-1* (CSHL_GT19652; Sundaresan et al., 1995) and *pip2;1-2 pip2;2-3* were used for crossing. | **5.** This work |
| | **6.** *pip2;1-2 pip2;2-3 pip2;7-2* | **6.** *pip2;7-2* (CRISPR/Cas9 deletion of 220 bp (nucleotides 156-375) and two further CRISPR/Cas9-induced mutations; Schley et al., 2025) and *pip2;1-2 pip2;2-3* were used for crossing. | **6.** This work |
| | **7.** *pip2;1-2 pip2;2-3 pip2;4-1 pip2;6-3* | **7.** *pip2;4-1* (SM_3_20853, Tissier et al., 1999), *pip2;6-3* (SALK_092140, Alonso et al., 2003) and *pip2;1-2 pip2;2-3* were used for crossing. | **7.** This work |
| | **8.** *pip2;1-2 pip2;2-3 pip2;4-1 pip2;6-3 pip2;7-4* | **8.** *pip2;7-4* (CRISPR/Cas9 deletion confirmed by PCR and sequencing, indicating a deletion of 376 bp (nucleotides 156 to 531 relative to the ATG start codon) was generated in *pip2;1-2 pip2;2-3 pip2;4-1 pip2;6-3*. | **8.** Poitout et al., 2026; this work. |

**Table S1**. Mutant and transgenic lines (continued)

| Allele(s) | Mutant /Transgenic line | Specification | Reference |
|---|---|---|---|
| | **9.** *pip1;1-1* (HA-PIP1;1) | **9.** pPm42GW,3_HA-PIP1;1 (Figure S10; Table S4) construct introduced into *pip1;1-1* mutant. | **9.** This work |
| | **10.** *pip1;1-1 pip2;1-2 pip2;2-3* (HA-PIP1;2) | **10. A h**omozygous, single insertion transgenic line harboring HA-PIP1;1 in *pip1;1-1* was crossed with *pip1;1-1 pip2;1-2 pip2;2-3.* | **10.** This work |
| | **11.** *pip1;2-2* (HA-PIP1;2) | **11.** pPm42GW,3_HA-PIP1;2 (Figure S10; Table S4) construct introduced into *pip1;2-2* mutant by floral dipping. | **11.** This work |
| | **12.** *pip1;2-2 pip2;1-2 pip2;2-3* (HA-PIP1;2) | **12.** A homozygous, single insertion transgenic line harboring HA-PIP1;2 in *pip1;2-2* was crossed with *pip1;2-2 pip2;1-2 pip2;2-3.* | **12.** This work |
| | **13.** *pip1;2-2* (EGFP-PIP1;2) | **13.** pPm42GW,3_EGFP-PIP1;2 (Figure S10; Table S4) construct introduced into *pip1;2-2* mutant by floral dipping. | **13.** This work |
| | **14.** *pip1;2-2 pip2;1-2 pip2;2-3* (EGFP-PIP1;2) | **14.** A homozygous, single insertion transgenic line harboring EGFP-PIP1;2 in *pip1;2-2* was crossed with *pip1;2-2 pip2;1-2 pip2;2-3.* | **14.** This work |
| | **15.** *pip1;2-2 pip2;1-2 pip2;2-3 pip2;7-5* (EGFP-PIP1;2) | **15.** *pip2;7-5* (CRISPR/Cas9 deletion (nucleotides 155 to 426 relative to the ATG start codon) generated in *pip1;2-2 pip2;1-2 pip2;2-3* (EGFP-PIP1;2) | **15.** This work |

**Table S2**. Plasmids.

| Serial no. | Plasmid | Description and Reference |
|---|---|---|
| 1. | pDGE332, pDGE333, pDGE335, pDGE337 | Shuttle vectors used for loading the *DLN1* sgRNAs (Ordon et al., 2017) (Table S4) |
| 2. | pDONR221-P1P4$^{TM}$ | Donor vector for GATEWAY$^{TM}$ cloning (ThermoFisher, Invitrogen, Germany). |
| 3. | pDONR221-P3P2$^{TM}$ | Donor vector for GATEWAY$^{TM}$ cloning (ThermoFisher, Invitrogen, Germany). |
| 4. | pDONR221-P4P1R$^{TM}$ | Donor vector for GATEWAY$^{TM}$ cloning (ThermoFisher, Invitrogen, Germany). |
| 5. | pDONR221$^{TM}$ | Donor vector for GATEWAY$^{TM}$ cloning (ThermoFisher, Invitrogen, Germany). |
| 6. | pPm42GW-At2S3$_{pro}$::GFP | At2S3$_{pro}$::GFPpA cassette amplified from pAlligator2 (Bensmihen et al., 2004) with flanking *Sac*I sites and cloned into *Sac*I site of pPm42GW,3 (Karimi et al., 2007) |
| 7. | pDGE347_*DLN1*_4X CRISPR/Cas9 construct | Plant transformation vector with Cas9 expression cassette and RFP for selection of transformants (Stuttmann et al., 2021). This was used to introduce mutations into *DLN1* gene at two target sites (Table S4); both target sites were introduced twice with the order 1-1-2-2. |
| 8. | pDGE250_*PIP2;7*_4X CRISPR/Cas9 construct | Schley et al. (2025) |
| 9. | pBiFCt-2in1-NN | 2in1 Destination vector for GATEWAY$^{TM}$ cloning (Grefen & Blatt, 2012). |
| 10. | pBiFCt-2in1-NN PIP1;1-YFP$^{N}$/PIP2;1-YFP$^{C}$ | *PIP1;1* ORF cloned into pDONR221-P3P2 and *PIP2;1* ORF cloned into pDONR221-P1P4 were used to generate the final BiFC expression vector by LR recombination reaction (Figure S9). |
| 11. | pBiFCt-2in1-NNPIP1;1-YFP$^{N}$/PIP2;2-YFP$^{C}$ | *PIP1;1* ORF cloned into pDONR221-P3P2 and *PIP2;2* ORF cloned into pDONR221-P1P4 were used to generate the final BiFC expression vector by LR recombination reaction (Figure S9). |
| 12. | pBiFCt-2in1-NNPIP1;1-YFP$^{N}$/AHA1-YFP$^{C}$ | *PIP1;1* ORF cloned into pDONR221-P3P2 and *AHA1* ORF cloned into pDONR221-P1P4 were used to generate the final BiFC expression vector by LR recombination reaction (Figure S9). |
| 13. | pBiFCt-2in1-NN PIP1;1-YFP$^{N}$/PIP2;7-YFP$^{C}$ | *PIP1;1* ORF cloned into pDONR221-P3P2 and *PIP2;7* ORF cloned into pDONR221-P1P4 were used to generate the final BiFC expression vector by LR recombination reaction (Figure S9). |
| 14. | pBiFCt-2in1-NN PIP1;2-YFP$^{N}$/DLN1-YFP$^{C}$ | *PIP1;2* ORF cloned into pDONR221-P3P2 and *DLN1* ORF cloned into pDONR221-P1P4 were used to generate the final BiFC expression vector by LR recombination reaction (Figure S9). |
| 15. | pBGWFS7 | Karimi et al. (2007); used as template to amplify EGFP, e.g., PCR2 component (Figure S10a) |
| 16. | pEN-R2-3xHA-L3 | Karimi et al. (2007), https://vectorvault.vib.be/collection/pen-r2-3xha-l3; used as template to amplify 3xHA-tag (HA), e.g., PCR2 component (Figure S10a) |

**Table S2**. Plasmids (continued)

| Serial no. | Plasmid | Description |
|---|---|---|
| 15. | pBiFCt-2in1-NN PIP1;2-YFP$^{N}$/HRD3A-YFP$^{C}$ | *PIP1;2* ORF cloned into pDONR221-P3P2 and *HRD3A* ORF cloned into pDONR221-P1P4 were used to generate the final BiFC expression construct by LR recombination reaction (Figure S9; Table S4). |
| 16. | pBiFCt-2in1-NN PIP1;2-YFP$^{N}$/CDC48A-YFP$^{C}$ | *PIP1;2* ORF cloned into pDONR221-P3P2 and *CDC48A* ORF cloned into pDONR221-P1P4 were used to generate the final BiFC expression construct by LR recombination reaction (Figure S9, Table S4). |
| 17. | pBiFCt-2in1-NN PIP1;2-YFP$^{N}$/ RMA1-YFP$^{C}$ | *PIP1;2* ORF cloned into pDONR221-P3P2 and *RMA1* ORF cloned into pDONR221-P1P4 were used to generate the final BiFC expression construct by LR recombination reaction (Figure S9, Table S4). |
| 18. | pBiFCt-2in1-NN PIP1;2-YFP$^{N}$/HRD1B-YFP$^{C}$ | *PIP1;2* ORF cloned into pDONR221-P3P2 and *HRD1B* ORF cloned into pDONR221-P1P4 were used to generate the final BiFC expression construct by LR recombination reaction.(Figure S9, Table S4). |
| 19. | pBiFCt-2in1-NN HRD3A-YFP$^{N}$/DLN1-YFP$^{C}$ | *HRD3A* ORF cloned into pDONR221-P3P2 and *DERLIN1* ORF cloned into pDONR221-P1P4 were used to generate the final BiFC expression construct by LR recombination reaction (Figure S9, Table S4). |
| 20. | pPm42GW_PIP1;2$_{pro}$::EGFP-PIP1;2_At2S3$_{pro}$:: GFP | PIP1;2$_{pro}$::EGFP-PIP1;2 inserted into pPm42GW-At2S3$_{pro}$::GFP (serial no. 6) (Figure S10, Table S4). |
| 21. | pPm42GW_PIP1;1$_{pro}$::HA-PIP1;1_At2S3$_{pro}$:: GFP | PIP1;1$_{pro}$::HA-PIP1;1 inserted into pPm42GW-At2S3$_{pro}$::GFP (serial no. 6) (Figure S10, Table S4). |
| 22. | pPm42GW_PIP1;2$_{pro}$::HA-PIP1;2_At2S3$_{pro}$::GFP | PIP1;2$_{pro}$::HA-PIP1;2 inserted into pPm42GW-At2S3$_{pro}$::GFP (serial no. 6) (Figure S10, Table S4). |
| 23. | p35S$_{pro}$::PIP1;2-EGFP | 35S$_{pro}$::PIP1;2-EGFP inserted into pDON221P4P1R (Figure S11, Table S4), used for transient expression |
| 24. | pBS 2X35S$_{pro}$::HA-GW:tNOS | *2X35S$_{pro}$::HA-GW:tNOS* cassette of pAlligator2 (Bensmihen et al., 2004) cloned by restriction and ligation (*Hind*III, *Kpn*I) into pBluescript KS (+) (Invitrogen, USA) |
| 25. | pBS 2X35S$_{pro}$::HA-PIP2;1-tPIP2;1 | *PIP2;1* amplified from genomic DNA (Table S4), recombined into pBS 2X35S$_{pro}$::HA-GW:tNOS, used for transient expression |

**Table S3**. Oligonucleotides used for genotyping.

| Oligonucleotide name | Sequence | Purpose | Reference |
|---|---|---|---|
| ATG7_F | CAGCGTGATCTGTGAGAACTG | Primer to detect wild-type allele | This work |
| ATG7_R | TTCTTGGAGCTGGTACATTGG | Primer to detect wild-type and insertion allele | This work |
| bri1-9_dCAPS_F | TACGTTTATATCAAAAACGATGGG | Distinction of *bri1-9/BRI1* | Jin et al., 2007 |
| bri1-9_dCAPS_R | ATGTATCCAGACAACATGTTTTAA | Distinction of *bri1-9/BRI1* | Jin et al., 2007 |
| DLN1_F | CGTAGGCTTGTCTCTTGTGG | Primer to detect CRISPR/Cas9 knockout | This work |
| DLN1_R | ATGGCCCTTTCTCGAGTTGA | Primer to detect CRISPR/Cas9 knockout | This work |
| dSPM1_F | CTTATTTCAGTAAGAGTGTGGGGTT TTGG | Insertion specific primer for SM lines | Tissier et al., 1999 |
| HRD1A_F | CTTGAGCTTATCCGTGACCTG | Primer to detect wild-type allele | This work |
| HRD1A_R | TGCTACTGTGTTTGCAGATGG | Primer to detect wild-type allele | This work |
| HRD1B_F | AGTGGCATCATTTCTGCAAAC | Primer to detect wild-type allele | This work |
| HRD1B_R | GGAAGGGCTCAGGTGATTAAG | Primer to detect wild-type allele | This work |
| Lba1-mod | GGTTCACGTAGTGGGCCATC | Insertion specific primer for SALK lines | Alonso et al., 2003 |
| PIP2;1_SM_F | AACATATAACGTTGGCAAAAA | Primer to detect wild-type allele | Da Ines, 2008 |
| PIP2;1_SM_ R | TGGTTAAGACAGGGTTAGTCA | Primer to detect wild-type allele | Da Ines, 2008 |
| PIP2;2_SAIL_F | AAGTTATAGAAATGGCCAAAGAC | Primer to detect wild-type allele | Da Ines, 2008 |
| PIP2;2_SAIL_R | CTCAAACGTTGGCTGCACTTCTG | Primer to detect wild-type allele | Da Ines, 2008 |
| PIP2;4_SM_F | CAAAGCATTCCAAAGTTCTTA | Primer to detect wild-type allele | Da Ines, 2008 |
| PIP2;4_SM_R | GCCTTTTCGTTGTTGTAAATA | Primer to detect wild-type allele | Da Ines, 2008 |
| PIP2;6_SALK_F | CTATCTCATCTGGATCAGCTGGTT | Primer to detect wild-type allele | This work |
| PIP2;6_SALK_ R | TACACACAAACCTCCCCCACA | Primer to detect wild-type allele | This work |
| PIP2;7-1_ F | GCTGTGACTTTCGGTCTGTTC | Primer to detect wild-type allele | This work |
| PIP2;7-1_ R | AAACCAAAGGCAAACGATTAAC | Primer to detect wild-type allele | This work |
| PIP2;7_ F | AAACCCACCATGGAAAAGACTA | Primer to detect CRISPR/Cas9 knockout | This work |
| PIP2;7_ R | GACTTACGGGGATGTGAGAGTC | Primer to detect CRISPR/Cas9 knockout | This work |
| SAIL_L_F | TTCATAACCAATCTCGATACAC | Insertion specific primer for SAIL lines | McElver et al., 2001; Sessions et al., 2002 |
| SAIL_L_R | ACCGGAACGTGGGAGTCTC | Insertion specific primer for SAIL lines | McElver et al., 2001; Sessions et al., 2002 |

**Table S4**. Oligonucleotides used for cloning. Oligonucleotides related to GATEWAY™ cloning include GW in their names and indicate the respective attB recombination module (sequences indicated by small letters); four G residues at the 5'-end are added for efficient cloning according to manufacturer's recommendation. Additional nucleotides potentially deviating from the specific gene sequences were introduced if required to generate in-frame fusions. *A. thaliana* (Col) genomic DNA, cDNA, or plasmid DNA was used as template unless otherwise indicated. F, forward primer; R, reverse primer.

| Oligonucleotide name | Sequence | Purpose | Reference |
|---|---|---|---|
| **Gateway cloning specific oligonucleotides used for generating pBiFCt-2in1-NN constructs** | | | |
| AHA1_ORF_GW_B1_F | GGGGacaagtttgtacaaaaaagcaggctTAATGTCAGGTCTCGAAGATA | Primer used to amplify *AHA1* from cDNA to construct pDONR221-P1P4_AHA1 | This work |
| AHA1_ORF_GW_B4_R | GGGGacaactttgtatagaaaagttgggtGCTACACAGTGTAGTGATGTC | Primer used to amplify *AHA1* from cDNA to construct pDONR221-P1P4_AHA1 | This work |
| CDC48A_ORF_GW_B1_F | GGGGacaagtttgtacaaaaaagcaggctTAATGTCTACCCCAGCTGAA | Primer used to amplify *CDC48A* from cDNA to construct pDONR221-P1P4_CDC48A | This work |
| CDC48A_ORF_GW_B4_R | GGGGacaactttgtatagaaaagttgggtCACTAATTGTAGAGATCATC | Primer used to amplify *CDC48A* from cDNA to construct pDONR221-P1P4_CDC48A | This work |
| DLN1_ORF_GW_B1_F | GGGGacaagtttgtacaaaaaagcaggctTAATGTCTTCTCCTGGCGA | Primer used to amplify *DLN1* from cDNA to construct pDONR221-P1P4_DLN1 | This work |
| DLN1_ORF_GW_B4_R | GGGGaccactttgtacaagaaagctgggtTTCAAACTTGTTACGACCAA | Primer used to amplify *DLN1* from cDNA to construct pDONR221-P1P4_DLN1 | This work |
| DLN1_ORF_GW_B3_F | GGGGacaactttgtataataaagttgGAATGTCTTCTCCTGGCGAATTCT | Primer used to amplify *DLN1* from cDNA to construct pDONR221-P3P2_DLN1 | This work |
| DLN1_ORF_GW_B2_R | GGGGaccactttgtacaagaaagctgggtTCAGTCGGTGAGACGATATGA | Primer used to amplify *DLN1* from cDNA to construct pDONR221-P3P2_DLN1 | This work |
| HRD1B_ORF_GW_B1_F | GGGGacaagtttgtacaaaaaagcaggctTAATGATTCAGCTAAAGGTTTACGC | Primer used to amplify *HRD1B* from cDNA to construct pDONR221-P1P4_HRD1B | This work |
| HRD1B_ORF_GW_B4_R | GGGGacaactttgtatagaaaagttgggtTCACCACTATGCAGTATCCG | Primer used to amplify *HRD1B* from cDNA to construct pDONR221-P1P4_HRD1B | This work |
| HRD3A_ORF_GW_B1_F | GGGGacaagtttgtacaaaaaagcaggctTAATGAGAATATTAAGCTACGG | Primer used to amplify *HRD3A* from cDNA to construct pDONR221-P1P4_HRD3A | This work |
| HRD3A_ORF_GW_B4_R | GGGGacaactttgtatagaaaagttgggtTTGATTACCGTGGGAACGCA | Primer used to amplify *HRD3A* from cDNA to construct pDONR221-P1P4_HRD3A | This work |
| HRD3A_ORF_GW_B3_F | GGGGacaactttgtataataaagttgGAATGAGAATATTAAGCTACGGAATCGT | Primer used to amplify *HRD3A* from cDNA to construct pDONR221-P3P2_HRD3A | This work |
| HRD3A_ORF_GW_B2_R | GGGGaccactttgtacaagaaagctgggtAATTGATTACCGTGGGAACGCAGC | Primer used to amplify *HRD3A* from cDNA to construct pDONR221-P3P2_HRD3A | This work |
| PIP1;1_ORF_GW_B3_F | GGGGacaactttgtataataaagttgTAATGGAAGGCAAGGAAGAAGACG | Primer used to amplify *PIP1;1* from cDNA to construct pDONR221-P3P2_PIP1;1 | This work |
| PIP1;1_ORF_GW_B2_R | GGGGaccactttgtacaagaaagctgggtTGCTTCTGGACTTGAAGGGGA | Primer used to amplify *PIP1;1* from cDNA to construct pDONR221-P3P2_PIP1;1 | This work |
| PIP1;2_ORF_GW_B3_F | GGGGacaactttgtataataaagttgTAATGGAAGGTAAAGAAGAAGATG | Primer used to amplify *PIP1;2* from cDNA to construct pDONR221-P3P2_PIP1;2 | This work |
| PIP1;2_ORF_GW_B2_R | GGGGaccactttgtacaagaaagctgggtTGCTTCTGGACTTGAATGGGA | Primer used to amplify *PIP1;2* from cDNA to construct pDONR221-P3P2_PIP1;2 | This work |
| PIP2;1_ORF_GW_B1_F | GGGGacaagtttgtacaaaaaagcaggctTAATGGCAAAGGATGTGGAAG | Primer used to amplify *PIP2;1* from cDNA to construct pDONR221-P1P4_PIP2;1 | This work |

**Table S4**. Oligonucleotides used for cloning (continued).

| Oligonucleotide name | Sequence | Purpose | Reference |
|---|---|---|---|
| PIP2;1_ORF_GW_B4_R | GGGGacaactttgtatagaaaagttgggtG GACGTTGGCAGCACTTCTGA | Primer used to amplify *PIP2;1* from cDNA to construct pDONR221-P1P4_PIP2;1 | This work |
| PIP2;2_ORF_GW_B1_F | GGGGacaagtttgtacaaaaaagcaggct TAATGGCCAAAGACGTGGAAG | Primer used to amplify *PIP2;2* from cDNA to construct pDONR221-P1P4_*PIP2;2* | This work |
| PIP2;2_ORF_GW_B4_R | GGGGacaactttgtatagaaaagttgggtG AACGTTGGCTGCACTTCTGA | Primer used to amplify *PIP2;2* from cDNA to construct pDONR221-P1P4_*PIP2;2* | This work |
| PIP2;7_ORF_GW_B1_F | GGGGacaagtttgtacaaaaaagcagg ctTAATGTCGAAAGAAGTGAGC GAAG | Primer used to construct pDONR221-P1P4_*PIP2;7* | This work |
| PIP2;7_ORF_GW_B4_R | GGGGacaactttgtatagaaaagttgggt AAAGCCTTCATTAATTGGTTGC | Primer used to construct pDONR221-P1P4_*PIP2;7* | This work |
| RMA1_ORF_GW_B1_ F | GGGGacaagtttgtacaaaaaagcaggct TGATGGCCTTAGATCAATCTT | Primer used to construct pDONR221-P1P4_*RMA1* | This work |
| RMA1_ORF_GW_B4_R | GGGGacaactttgtatagaaaagttgggt GAACAGTAGCAGAACCAAGC | Primer used to construct pDONR221-P1P4_*RMA1* | This work |
| **Oligonucleotides used for generating EGFP/HA-tagged PIP1;1 and PIP1;2 constructs using GATEWAY™ cloning** | | | |
| PIP1;1_Pro_GW_B4_F | GGGGacaactttgtatagaaaagttgAAA GCATGGTAAAATTGGTG | PCR1 forward (Figure S10) | This work |
| PIP1;1_Pro_GW_B1R_R | GGGGactgcttttttgtacaaacttgATCTT CGATCTCTGTAGAGAGAAAT | PCR1 reverse (Figure 10) | This work |
| EGFP_GW_B1_F | GGGGacaagtttgtacaaaaaagcaggct ATGGTGAGCAAGGGCG | PCR2 and PCR4 forward (Figure S10) | This work |
| PIP1;1_TER_GW_B2_R | GGGGaccactttgtacaagaaagctgggtC TCGTGGAATGATCAAACTT | PCR3 and PCR4 reverse (Figure S10) | This work |
| EGFP _PIP1;1_ORF_Fusion_F | catggacgagctgtacaagGGCGCTAT GGAAGGCAAGGAAGAAGAC | PCR3 forward with linker encoding GA in between EGFP and PIP (Figure S10) | This work |
| EGFP _PIP1;1_ORF_Fusion_R | GTCTTCTTCCTTGCCTTCCATAG CGCCcttgtacagctcgtccatg | PCR2 reverse with linker encoding GA in between EGFP and PIP (Figure S10) | This work |
| HA_GW_B1_F | GGGGacaagtttgtacaaaaaagcaggct ATGgcataccccttacgatg | PCR2 and PCR4 forward (Figure S10) | This work |
| HA_PIP1;1_ORF_Fusion_F | CGACGTTCCAGATTACGCTGGC GCTATGGAAGGCAAGGAAGAAG | PCR3 forward with linker encoding GA in between HA and PIP (Figure S10) | This work |
| HA_PIP1;1_ORF_Fusion_R | GTCTTCTTCCTTGCCTTC**CAT**AG CGCCAGCGTAATCTGGAACGTC G | PCR2 reverse with linker encoding GA in between HA and PIP (Figure S10) | This work |
| PIP1;2_Pro_GW_B4_F | GGGGacaactttgtatagaaaagttgTCG AATCTTCCTCATTTGAA | PCR1 forward (Figure S10) | This work |
| PIP1;2_Pro_GW_B1R_R | GGGGACTGC**TTTTTT**GTACAAAC TTGctctctctctctttctctctagagc | PCR1 reverse (Figure S10) | This work |
| EGFP _PIP1;2_ORF_Fusion_R | ACATCTTCTTCTTTACCTTCCATA GCGCCcttgtacagctcgtccatg | PCR2 reverse with linker encoding GA in between EGFP and PIP (Figure S10) | This work |
| EGFP _PIP1;2_ORF_Fusion_F | catggacgagctgtacaagGGCGCTAT GGAAGGTAAAGAAGAAGATGT | PCR3 forward with linker encoding GA in between EGFP and PIP (Figure S10) | This work |

**Table S4**. Oligonucleotides used for cloning (continued).

| Oligonucleotide name | Sequence | Purpose | Reference |
|---|---|---|---|
| PIP1;2_TER_GW_B2_R | GGGGACCAC**TTTGTA**CAAGAAAGCTGGGTatgccttggtaattcagaca | PCR3 reverse (Figure S10) | This work |
| HA_PIP1;2_ORF_Fusion_R | ACATCTTCTTCTTTACCTTCCATAGCGCCAGCGTAATCTGGAACGTCG | PCR2 reverse (Figure S10) | This work |
| HA_PIP1;2_ORF_Fusion_F | cgacgttccagattacgctGGCGCTATGGAAGGTAAAGAAGAAGATGT | PCR3 forward (Figure S10) | This work |
| 35S_GW_B4_F | GGGGacaactttgtata**gaaaa**gttgATTTAGGTGACACTATAGAATACTCAAG | PCR1 and PCR4 forward to amplify CaMV 35Spro module and final fusion construct for generating $35S_{pro}$::PIP1;2-EGFP recombined into pDONR-P4P1R (Table S2, S4; Figure S11) | This work |
| PIP1;2_ORF_35S _ Fusion_R | ACATCTTCTTCTTTACCTTCCATCGACTAGAATAGTAAATTGTAATGTTG | PCR1 reverse to amplify 35Spro module for generating $35S_{pro}$::PIP1;2-EGFP (Table S2, S4; Figure S11) | This work |
| PIP1;2_ORF_35S _ Fusion_F | CAACATTACAATTTACTATTCTAGTCGATGGAAGGTAAAGAAGAAGATGT | PCR2 forward to amplify PIP1;2-ORF module for generating $35S_{pro}$::PIP1;2-EGFP (Table S2, S4; Figure S11) | This work |
| PIP1;2_ORF_EGFP_ Fusion_R | CGCCCTTGCTCACCATAGCGCCGCTTCTGGACTTGAATGG | PCR2 reverse to amplify PIP1;2-ORF module with C-terminal overlap with EGFP and a GA-encoding linker in between for generating $35S_{pro}$::PIP1;2-EGFP (Table S2, S4; Figure S11) | This work |
| PIP1;2_ORF_EGFP_ Fusion_F | CCATTCAAGTCCAGAAGCGGCGCTatggtgagcaagggcg | PCR3 forward to amplify EGFP module for generating $35S_{pro}$::PIP1;2-EGFP fusion (Table S2, S4; Figure S11) | This work |
| EGFP_STOP_GW_B1R_R | GGGGACTGCTTTTTTGTACAAACTTGttacttgtacagctcgtccat | PCR3 and PCR4 reverse to amplify EGFP module and final fragment $35S_{pro}$::PIP1;2-EGFP into pDONRP4P1R (Table S2, S4; Figure S11) | This work |
| PIP2;1_ORF_GW_B1_F | GGGGACAAGTTTGTACAAAAAAGCAGGCTccatggcaaaggatgtggaagc | Amplification of PIP2;1 cDNA for recombination into pBS $2X35S_{pro}$::HA-GW:tNOS | This work |
| PIP2;1_ORF_GW_B2_R | GGGGACCACTTTGTACAAGAAAGCTGGGTttagacgttggcagcacttc | Amplification of PIP2;1 cDNA for recombination into pBS $2X35S_{pro}$::HA-GW:tNOS | This work |

**Table S4.** Oligonucleotides used for cloning (continued).

| Oligonucleotide name | Sequence | Purpose | Reference |
|---|---|---|---|
| **CRISPR/Cas9 specific oligonucleotides used for generating CRISPR/Cas9 constructs** | | | |
| *DLN1*_gR1_F | ATTGAATTGATAGAGAAACCACCG | oligonucleotide used for generating the single guide RNA for pDGE347_*DLN1*_4X CRISPR/Cas9 construct | This work |
| *DLN1*_gR1_R | AAACCGGTGGTTTCTCTATCAATT | oligonucleotide used for generating the single guide RNA for pDGE347_*DLN1*_4X CRISPR/Cas9 construct | This work |
| *DLN1*_gR2_F | ATTGAGCAACTAAGCCAAGCTGCG | oligonucleotide used for generating the single guide RNA for pDGE347_*DLN1*_4X CRISPR/Cas9 construct | This work |
| *DLN1*_gR2_R | AAACCGCAGCTTGGCTTAGTTGCT | oligonucleotide used for generating the single guide RNA for pDGE347_*DLN1*_4X CRISPR/Cas9 construct | This work |

**Table S5**. Oligonucleotides used for RT-qPCR.

| Oligonucleotide name | Forward primer | Reverse primer | Reference |
|---|---|---|---|
| *BiP 1/2* AT5G28540 *BiP1*& AT5G42020 (*BiP2*) | TACGTGTACAACATGAAG AACC | TCTTCTTTCTCTGAG TTTTGGT | Designed by Primer 3 (Untergasser et al., 2012) |
| *BiP 3* AT1G09080 | ACAAGCTTGAAACGTATG TGTA | CACATTCTCTTCTAA CCACTCC | Designed by Primer 3 (Untergasser et al., 2012) |
| *DLN1* AT4G29330 | AAACAATGCCACTAACTA ATCC | TAGAAACATGAACCA CATTTTG | Designed by Primer 3 (Untergasser et al., 2012) |
| *EF1ALPHA* AT5G60390 | GAGCACGCTCTTCTTGCT TTCA | TCAGGTATGAAGAC ACCTCCTT | Amplicon 133 bp, Bridging Intron, Czechowski *et al.,* (2005) and own design |
| *GRF1* AT4G09000 | TCATCGATCGAACAAAAG GA | AAGCTTAAGGATTCC GTCACA | Designed by Primer 3 (Untergasser et al., 2012) |
| *HRD1B* AT1G65040 | GGTACTTCAAGTTCTGAT GGTC | GATAAACAATGGATC TCCCATA | Designed by Primer 3 (Untergasser et al., 2012) |
| *PIP1;1* AT3G61430 | CTGGCCTTGTCCTTAGTT GCTTC | TCTCCTTTGGAACTT CTTCCTTG | Postaire et al., 2010 |
| *PIP1;2* AT2G45960 | TCCTCTTCTTTGCCTAAT GGAGAC | AGTTGCCTGCTTGA GATAAAC | Postaire et al., 2010 |
| *PIP1;3* AT1G01620 | GCTGTGGATGATCTGGT TTTATCG | GCCGAAACAATATG GATCTTACTC | Postaire et al., 2010 |
| *PIP1;4* AT4G00430 | CTCTGAAGTCTAAGGTGA TTAGTGC | CAACCCGAGAACTT GATGTTGA | Postaire et al., 2010 |
| *PIP1;5* AT4G23400 | TGTTTCCTATGTCATGTG TGATG | GTACACAATGTATTC TTCCATTGAC | Postaire et al., 2010 |
| *RHIP 1* AT4G26410 | GAGCTGAAGTGGCTTCC ATGAC | AAATTGTGGATTTGT GTTGGAT | Amplicon 123 bp, bridging intron; Selected by Czechowski *et al.,* 2005 and Souček *et al.,* 2017 and own design |
| *S16* AT5G18380 | TCTGGTAACGAGAACGA GCAC | TTTACGCCATCCGTC AGAGTAT | Designed by Primer3 (Untergasser et al., 2012) |
| *TUBULIN6* AT5G12250 | CCCTCGTCTCCACTTTTT | GTTCTTTGAATCCCA CATCT | Souček *et al.,* 2017 |
| *UBIQUITIN5* AT3G62250 | GATGGATCTGGAAAGGT TCAG | ATCTACCGCTACAAC AGATCAAG | Designed by Primer3 (Untergasser et al., 2012) |
